# Upstream Stimulatory Factors regulate HIV-1 latency and are required for robust T cell activation

**DOI:** 10.1101/2023.04.21.537777

**Authors:** Riley M. Horvath, Ivan Sadowski

**Author notes:** Correspondence to: I. Sadowski, Dept. of Biochemistry and Molecular Biology, UBC, 2350 Health Sciences Mall, Vancouver, B.C., V6T 1Z3, CANADA.

## Abstract

RBF-2, composed minimally of the factors USF1, USF2, and TFII-I, is the cognate binding complex of stringently conserved *cis*-elements on the HIV-1 LTR, designated RBE3 and RBE1. Mutations of these elements prevent induction of provirus in response to T cell signaling. However, the function of USF1 and USF2 for this effect are relatively uncharacterized. Here, we find that deletion of the *USF2* but not *USF1* gene in T cells abrogates HIV-1 expression. Loss of *USF2* caused a reduction in expression of USF1 protein, an effect that was not associated with decreased *USF1* mRNA abundance. USF1 and USF2 were previously shown to exist predominately as heterodimers on DNA *in vivo* and cooperatively regulate target genes. To examine this, we performed RNA-seq analysis of T cell lines bearing knockouts of genes encoding these factors. In untreated cells we found limited evidence of coordinated global gene regulation between USF1 and USF2. In contrast we observed a high degree of genome-wide cooperative regulation of RNA expression between these factors in cells stimulated with the combination of PMA and ionomycin. In particular, we found that *USF1* deletion resulted in a restricted T cell activation response, and this phenotype was exaggerated in *USF2* knockout cells. These observations indicate that USF2, but not USF1, is crucial for HIV-1 expression, but the combined function of these factors is required for full T cell response.

## Introduction

Despite decades of research, the etiological agent of acquired immunodeficiency syndrome (AIDS), human immunodeficiency virus (HIV-1), remains a pressing global healthcare issue as nearly 40 million individuals are infected globally and close to a million deaths occur annually (UNAIDS 2022). Current anti-retroviral therapy (ART) typically limits HIV-1 replication in most individuals to a chronic symptomless infection, so long as treatment is maintained (Temereanca and Ruta 2023). However, this treatment must be maintained for the lifespan of infected individuals to prevent rebound of viral replication from latently infected cells (Sadowski and Hashemi 2019), and often altered because of the emergence of drug resistant HIV-1 variants (Puertas et al. 2020). The barrier to a cure for HIV-1/ AIDS infection is the population of infected CD4^+^ T memory cells that establish proviral latency early upon infection. This extremely long-lived latently infected cell population provides a reservoir of viremia that evades host immune surveillance and is invulnerable to ART (Finzi 1997)(Joos et al. 2008). Various proposed strategies directed towards eliminating latently infected cells involve modulating expression of latent provirus (Sadowski and Hashemi 2019). Often referred to as “shock and kill” and “block and lock”, these strategies would employ latency reversing and latency promoting agents, respectively, and have been the subject of intense recent investigation (Abner and Jordan 2019)(Pasquereau and Herbein 2022). However, initial trials using latency reversing agents in patients on antiretroviral therapy did not produce significant reduction in the pool of latently infected CD4^+^ T cells *in vivo* (Sadowski and Hashemi 2019)(Petravic et al. 2017). Consequently, it is recognized that treatment with latency modulating agents must be applied with consideration of effects on global immune response, including function of CD8^+^ T cells (Walker-Sperling et al. 2016)(Y. Kim, Anderson, and Lewin 2018). Overall, however, development of effective therapies involving modulation of HIV-1 provirus expression will require a more detailed understating of mechanisms regulation HIV-1 latency.

The 5’ long terminal repeat (LTR) serves as the HIV-1 promoter and enhancer and possesses numerous *cis*-elements that bind host cell transcription factors (Pereira 2000)(Sadowski, Lourenco, and Malcolm 2008). Multiple *cis*-elements within the 5’ LTR bind transcription factors regulated downstream of T cell receptor signaling, causing provirus expression to be tightly linked to CD4 receptor engagement (Brooks et al. 2003). The Ras- response factor binding elements 3 and 1 (RBE3 and RBE1) are highly conserved on HIV-1 provirus in patients who develop AIDS (Mario Clemente Estable et al. 1996), and were initially characterized as required for induction of the HIV-1 LTR to activated Ras and MAPK signaling (Bell and Sadowski 1996). These conserved elements bind a transcription factor termed RBF-2, comprised minimally of the TFII-I, USF1 and USF2 proteins (Chen et al. 2005). This complex was found to be associated with the factor Yin Yang 1 (YY1) at the upstream RBE3 element which plays a role in establishing and enforcing latency (Bernhard et al. 2013). Mutations of the RBE1 and RBE3 elements abrogate binding of RBF-2 (USF1/2, TFII-I) *in vivo*, and render proviruses incapable of transcriptional induction in response to T cell signaling (Chen et al. 2005) (Malcolm et al. 2007) (Malcolm et al. 2008) (Matthew S. Dahabieh et al. 2011). This effect is mediated at least partially by recruitment of the co-factor TRIM24 to the HIV-1 LTR resulting in stimulated transcriptional elongation (Horvath, Brumme, and Sadowski 2023)(Horvath et al. 2023). However, despite that USF1 and USF2 were among the first cellular factors shown to bind the HIV-1 LTR (Roy 1997)(Du, Roy, and Roeder 1993), the functional significance of these factors for regulation of provirus transcription and latency are unknown.

Upstream Stimulatory Factor (USF) was initially identified as a transcriptional regulator which binds an E-box of the adenovirus major late promoter (MLP) (Sawadogo 1985). This factor is comprised of a heterodimer of two structurally related helix-loop-helix leucine zipper (b-HLH-LZ) proteins of 43 kDa and 44 kDa, designated USF1 and USF2, respectively (Sawadogo et al. 1988), expressed from individual genes located on separate chromosomes (Shieh et al. 1993) (Groenen et al. 1996). These factors are ubiquitously expressed, although relative ratios in abundance of the two proteins vary depending on tissue type (Gregor, Sawadogo, and Roeder 1990) (Sirito et al. 1994). USF1 and 2 have overall sequence identity of 44% with the C-terminal b-HLH-LZ displaying ∼70% conservation (Sirito et al. 1994). Given the high degree of sequence homology, similar effects for binding and activation of the Adenovirus MLP (Sawadogo et al. 1988), and predominance of USF1/USF2 heterodimers within cells (Viollet et al. 1996), functional differences between these two factors have largely been overlooked. However, *USF1* and *USF2* knockout mice display unique phenotypes, likely representing differential gene expression (Sirito et al. 1998), indicating these factors have distinct functions.

As for other members of the b-HLH transcription factor family, USFs have been observed to bind E-Box elements possessing the core CANNTG sequence, including within the Adenovirus MLP and HIV-1 RBE1 (CAGCTG) (Baxevanis and Vinson 1993) (Rada-Iglesias et al. 2008) (Matthew S. Dahabieh et al. 2011). However, USF binding is not limited to E-Box elements, as these factors also bind pyrimidine rich initiator (Inr) elements and the non-canonical E-box RBE3 element of HIV-1 in association with TFII-I (Roy 1997) (Chen et al. 2005) (Malcolm et al. 2008). USF1 and 2 are associated with both positive and negative effects on transcription, effects that are dependent upon target promoter context and cellular differentiation state (Andrews 2001) (Qyang et al. 1999). The temporal and spatial nature of USF function produce differential effects of these factors in specific cell types. For example, USF1 inhibits inflammatory NFκB signaling in lung tissue (Tiruppathi et al. 2014), while USF2 activity correlates with expression of proinflammatory cytokines in Th17 cells and refractory rheumatoid arthritis (Hu et al. 2020). Furthermore, USF1 plays an important role in UV response and prevention of tumor development (Galibert 2001) (Corre et al. 2004, 204)(Bouafia et al. 2014), while both USF1 and USF2 activate HOXA9 expression and have a driver role in leukemia (Zhang et al. 2020).

Although mechanism(s) are ill-defined, regulation of transcription by USF has been reported to involve alterations to chromatin organization. The HS4 insulator element of the chicken β*- globin* locus requires USF binding to retain H3K4me and H3ac and prevent encroachment of heterochromatin (West et al. 2004). Additionally, activation of lipogenic genes in response to insulin involves USF1 mediated recruitment of BAF60c and subsequent chromatin remodeling (Wang et al. 2013). The USFs have been shown to be regulated by signaling pathways through phosphorylation (Horbach et al. 2015) which regulates both DNA binding function and protein-protein interactions (Nowak et al. 2005) (Spohrer et al. 2017). Phosphorylation of USF2 by CDK5 is thought to contribute to carcinogenesis (Chi et al. 2019) while CK2 dependent phosphorylation of USF1 causes altered expression of genes involved in metabolism (Lupp et al. 2014) (Spohrer et al. 2017). Moreover, T cell activation induced by the phorbol-ester PMA causes extensive phosphorylation of both USF1 and USF2, although the regulatory implications of these modifications have not been determined (Chen et al. 2005).

Here, we demonstrate that HIV-1 provirus expression is inhibited upon depletion of *USF2,* whereas loss of *USF1* has no significant effect. Additionally, loss of USF2 results in a reduction of USF1 protein expression, an effect that does not correlate with *USF1* mRNA transcript abundance. Furthermore, we found that USF1 and USF2 are key factors required for T cell activation, as loss of either protein caused global constraint of T cell signal induced genes. *USF2* gene knockout limited T cell activation to a greater extent than a *USF1* knockout, and rendered T cells more restrictive to HIV-1 infection. These observations clarify the role of USF1 and USF2 for regulation of HIV-1 transcription, and identify these proteins as key signal regulated factors that play a vital role in activation of the T cell inflammatory response.

## Results

### Regulation of HIV-1 expression by USF1 and USF2

Previous studies have implicated USF1 and/or USF2 as regulators of HIV-1 expression, mostly based on transient transfections assays and LTR mutations that abolish binding of these factors (Du, Roy, and Roeder 1993) (d’Adda di Fagagna et al. 1995) (Sieweke 1998)(Chen et al. 2005)(Matthew S. Dahabieh et al. 2011). To examine specific effects of USF1 and USF2 on HIV-1 expression we produced *USF1* and *USF2* gene knockouts (KO) (Fig. 1*A*, 1*B*) in the Jurkat Tat mHIV-Luc T cell line bearing an integrated HIV-1 mini-virus where luciferase expression is driven by the 5’ LTR (Bernhard et al. 2011)(Horvath et al. 2023)(Horvath, Brumme, and Sadowski 2023). Surprisingly, we noted that USF1 protein abundance was reduced in all *USF2* knockout lines (Fig. 1*B*), whereas in contrast USF2 protein expression was not affected by *USF1* knockout (Fig. 1*A*). We examined activation of HIV-1 expression in these knockout lines and found that *USF1* disruption caused a slight increase in luciferase expression in most *USF1* knockout clones in unstimulated as well as T cell activating conditions as mediated by the phorbol ester PMA, which partially mimics stimulation of cells by T cell receptor engagement (Fig. 1*C*). In contrast, provirus expression was diminished in both unstimulated and stimulated conditions in all *USF2* knockout lines (Fig. 1*D*). Consistent with results using the knockout lines, we found that that shRNA mediated depletion of USF1 (Fig. S1*A*) had no effect on HIV-1 provirus expression (Fig. S1*B*), while shRNA knockdown of USF2 (Fig. S1*C*) inhibited HIV-1 expression in both untreated cells and cells stimulated with PMA (Fig. S1*D*).

**Figure 1.**
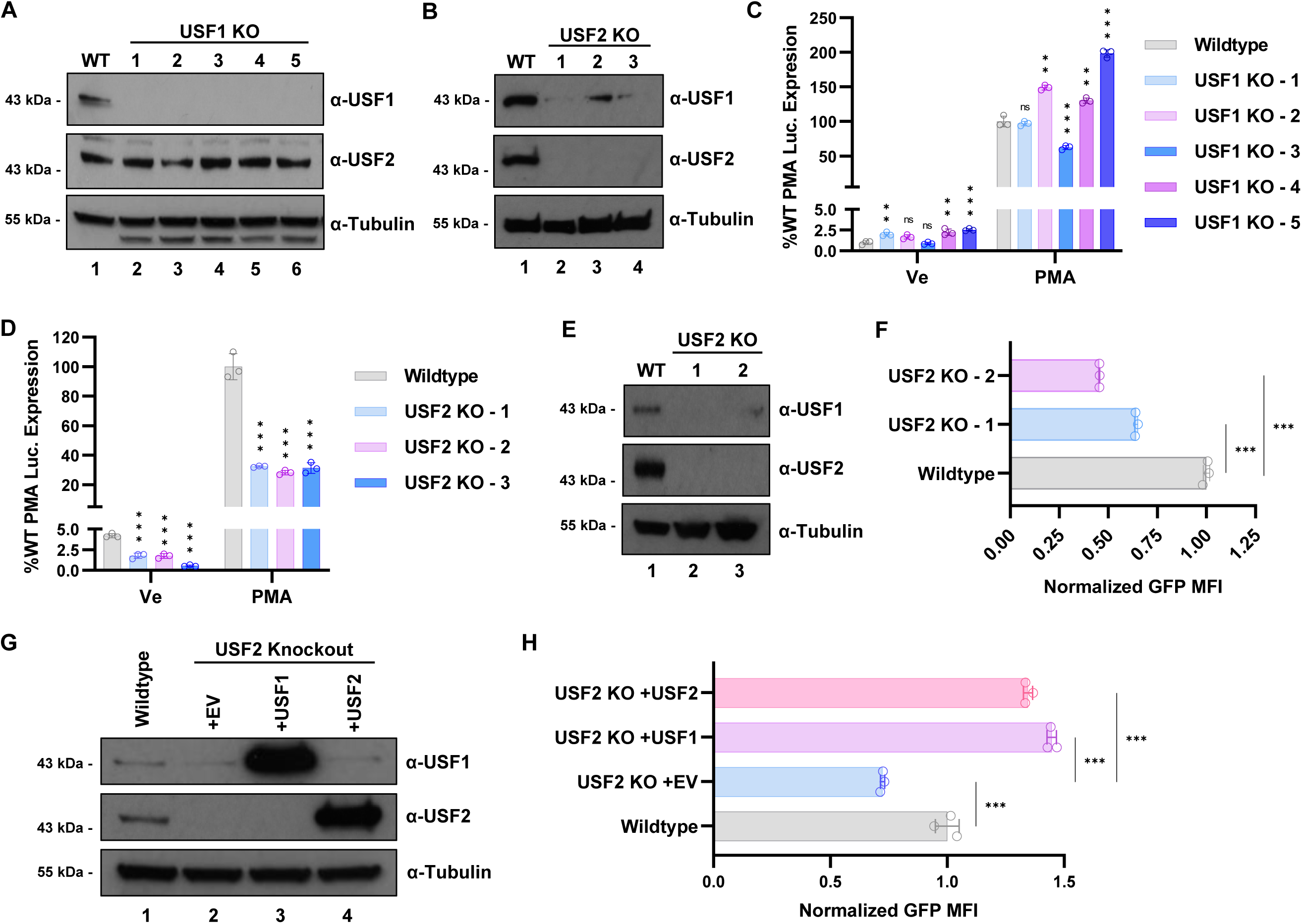
*USF2* knockout inhibits HIV-1 expression. **Panels A, B:** *USF1* (A) and *USF2* (B) KO cell lines were produced using CRISPR-Cas9 in the Jurkat mHIV-Luciferase parental cell line. I mmunoblotting of cellular lysates was performed using antibodies against USF1, USF2, and Tubulin. **Panel C:** Wildtype or *USF1* KO mHIV-Luciferase cells were left untreated (Ve, DMSO) or incubated with 5 nM PMA for 4 hours prior to measuring luciferase activity (*n* = 3, mean ±LSD). **Panel D:** Same as in C but *USF2* KO cell lines were analysed (*n* = 3, mean ±LSD). **Panel E:** *USF2* KO clonal cell lines were generated from the HEK293T parental line through CRISPR-Cas9 gene editing. KO was confirmed by immunoblotting whole cell extracts using antibodies against USF1, USF2, and Tubulin. **Panel F:** Wildtype and *USF2* KO HEK293T cells were co-transfected with LAI derived LTR-Tat-IRES-GFP reporter plasmid and EF1α-RFP expression vector (Fig. S2*A*). The GFP mean fluorescent intensity (MFI) of the RFP expressing population was determined one day post-transfection by flow cytometry and normalized to wildtype cells (*n* = 3, mean ±LSD). **Panel G:** HEK293T *USF2* KO cells were transfected with an empty vector (EV) or vectors expressing either USF1 or USF2. Whole cell lysates were examined by immunoblotting with antibodies against USF1, USF2, and Tubulin. **Panel H:** Wildtype and *USF2* KO HEK293T cell lines were co-transfected with LAI LTR-Tat-IRES-GFP reporter plasmid and either CMV-EV-EF1α-RFP (EV), CMV-USF1-EF1α-RFP (USF1) or CMV-USF2-EF1α-RFP (USF2) vector (Fig. S2*A*). One day post-transfection, the GFP mean fluorescent intensity (MFI) of the RFP+ population was determined by flow cytometry and normalized to CMV-EV-EF1α-RFP (EV) transfected wildtype cells (*n* = 3, mean ±LSD).

As USF1 protein expression was dependent upon USF2 in Jurkat T cells we examined the effect of *USF2* gene knockout in HEK293T cells, where we also observed loss of both USF1 and USF2 protein expression (Fig. 1*E*). We examined the effect of *USF2* knockout in these cells using transfection with an LAI-derived LTR-Tat-IRES-GFP reporter construct (Fig. S2*A*, S2*B*). Consistent with the above results, we observed significantly lower expression of GFP expression in HEK293T *USF2* knockout lines transfected with this reporter relative to WT (Fig. 1*F*). Remarkably, we found that co-transfection of HEK293T *USF2* KO cells with vectors expressing either USF1 or USF2 (Fig. 1*G*) with the HIV-1 LTR-Tat-IRES-GFP reporter not only rescued LTR transcription, but caused activation of GFP expression greater than that of WT HEK293T cells (Fig. 1*H*, S2*C*). These observations suggest that any combination of USF1 and USF2 can activate transcription from the HIV-1 LTR, but because USF1 protein expression is significantly decreased in *USF2* knockout lines while USF2 expression is not affected in *USF1* knockouts, only the *USF2* knockouts produce a significant effect on HIV-1 LTR transcription in T cells.

### Effect of USF1 and 2 on response of HIV-1 to latency reversing agents and Tat

We examined the effect of *USF1* and *USF2* knockouts on reactivation of HIV-1 in response to various mechanistically diverse latency reversing agents (LRAs). In T cells, the phorbol ester PMA activates protein kinase C (PKC) and Ras-MAPK signaling, while ionomycin causes release of intracellular calcium activating calcineurin, and treatment with this combination mimics T cell activation produced by CD4 receptor engagement (Sadowski and Mitchell 2005). We found that *USF2* KO inhibited HIV-1 provirus activation in response to PMA or ionomycin individually, or in combination (Fig. 2*A*). In contrast USF1 KO caused a slight increase in response to the combination of PMA/ionomycin or PMA alone (Fig. 2*A*). The effects of the *USF1* and *USF2* knockouts on response to the combination of PMA and ionomycin was most pronounced at 6 hours post treatment (Fig. 2*B*). As observed previously, we found that HIV-1 expression becomes repressed upon prolonged treatment (20 hrs) with PMA/ ionomycin in WT cells (Malcolm et al. 2007), and this effect was not significantly different in the *USF1* and *USF2* knockout lines (Fig. 2B). Interestingly, loss of USF1 caused slightly decreased induction following treatment with ionomycin alone (Fig. 2*A*), indicating that USF1 is not required for HIV-1 reactivation in response to PKC and Ras-MAPK signaling but may play a role for activation by calcium-calcineurin signaling. Ingenol 3-angelate (PEP005) also activates PKC signaling (Kedei et al. 2004), and similar to the results with PMA, we observed a significantly lower response to PEP005 in *USF2* KO cells, but no effect in the *USF1* KO line (Fig. 2*A*). These results indicate that USF2 is required for reactivation of HIV-1 in response to PKC, Ras-MAPK, and calcineurin T cell signaling pathways, whereas USF1 may cause some inhibition of response to MAPK activation.

**Figure 2.**
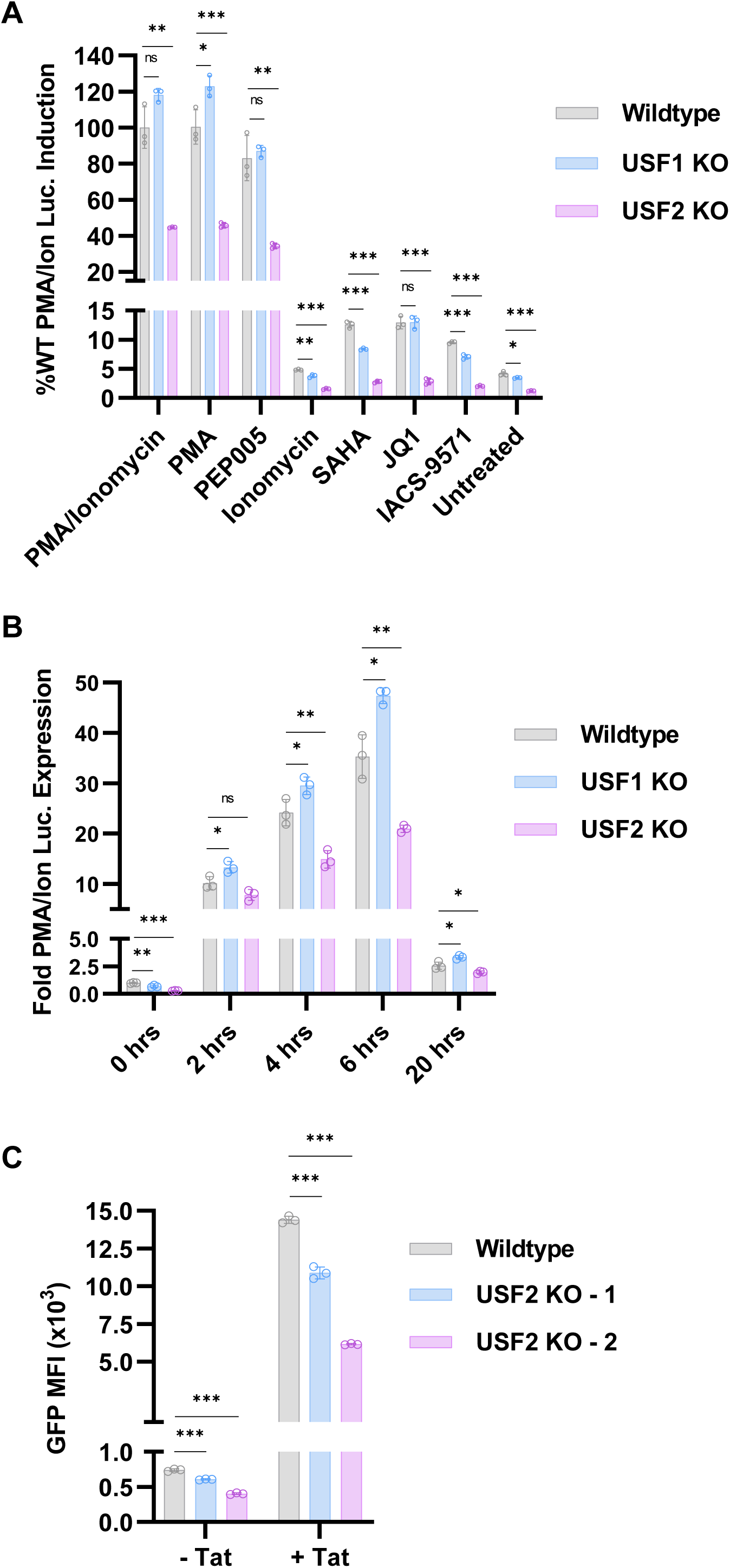
Effect of USF1 and USF2 on induction of HIV-1 in response to latency reversal agents and Tat. **Panel A:** Wildtype, *USF1* KO, or *USF2* KO mHIV-Luciferase cells were left untreated or treated with 5 nM PMA/ 1 μM ionomycin, 5 nM PMA, 10 nM PEP005, 1 μM ionomycin, 10 μM SAHA, 10 μM JQ1, or 10 μM IACS-9571 for 4 hrs prior to luciferase assay (*n* = 3, mean ±LSD). **Panel B:** Wildtype, *USF1* KO, or *USF2* KO mHIV-Luciferase were treated with 5 nM PMA and 1 μM ionomycin for the indicated amount of time, at which point luciferase expression was measured (*n* = 3, mean ±LSD). **Panel C:** Wildtype or *USF2* KO HEK293T cell lines were co-transfected with CMV-EV-EF1α-RFP vector and either an LTR- GFP (-Tat) or LTR-Tat-IRES-GFP (+Tat) reporter construct (Fig. S3*A*). One day post- transfection, the mean fluorescent intensity (MFI) of GFP for the RFP+ population was determined by flow cytometry (*n* = 3, mean ±LSD).

Latent provirus can be reactivated independent of T cell signaling by altering the epigenetic composition or interaction of bromodomain containing proteins with the LTR. Treatment with the histone deacetylase inhibitor (HDACi) suberanilohydroxamic acid (SAHA) (Desimio, Giuliani, and Doria 2017) modestly increased HIV-1 expression, an effect that was limited by both the *USF1* and *USF2* knockouts (Fig. 2*A*), indicating that these factors regulate HIV-1 latency through organization of LTR chromatin structure involving histone acetylation. Next, we examined effect of the USFs on reactivation in response to the bromodomain inhibitors JQ1 and IACS-9571. JQ1 is a well-studied LRA that inhibits the BRD4 bromodomain causing increased Tat transactivation (Li et al. 2013), while IACS-9571 inhibits the TRIM24 bromodomain causing increased LTR occupancy of this factor and stimulation of transcriptional elongation (Horvath et al. 2023; Horvath, Brumme, and Sadowski 2023) (Horvath et al, 2023). We found that the *USF2* knockout significantly limited the effect of JQ1, whereas *USF1* KO had no effect (Fig. 2A). In contrast, the *USF2* knockout, and to a lesser extent *USF1* KO inhibited reactivation of HIV-1 expression in cells treated with IACS-9571 (Fig. 2*A*), suggesting that both USFs impact regulation of HIV-1 by TRIM24. This is consistent with observations that TRIM24 is recruited to the LTR by RBF-2 (Horvath et al. 2023), which is comprised USF1, USF2, and TFII-I (Chen et al. 2005).

We also examined the effect of *USF2* gene knockout on activation of HIV-1 expression by the viral transactivator protein (Tat). For this experiment we transfected *USF2* knockout HEK293T lines (Fig. 1*E*) with HIV-1 LTR-Tat-IRES-GFP (+Tat), or LTR-GFP reporter plasmids (-Tat) (Fig. S3*A*, S3*B*). We observed that GFP expression from both HIV-1 LTR reporter plasmids was significantly lower in the *USF2* null cells compared to WT (Fig. 2*C*, S3*C*, S3*D*). These observations indicate that full activation of HIV-1 transcription by Tat requires USF2 protein, although USF2 also has a significant effect on HIV-1 expression in the absence of Tat.

**Figure 3.**
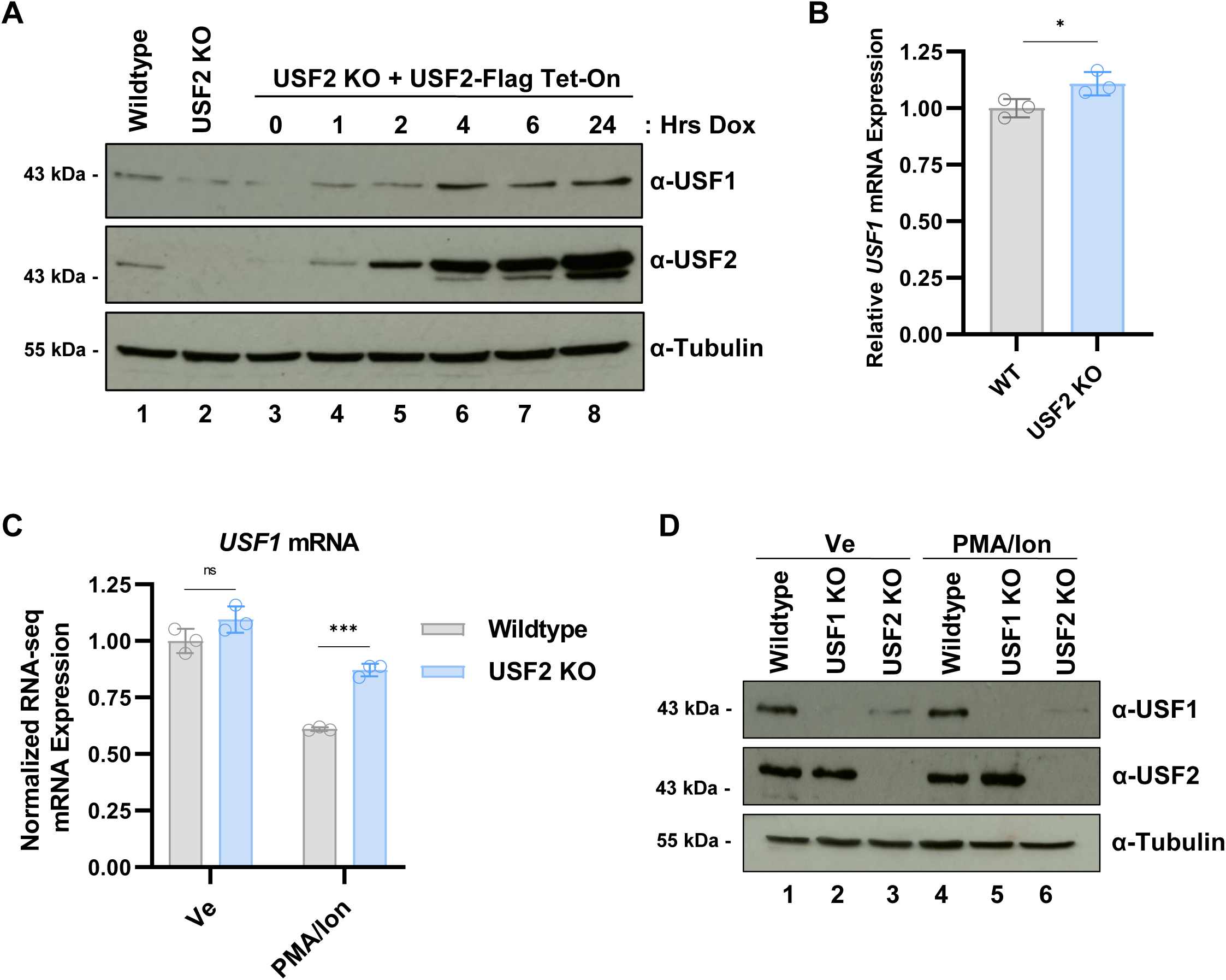
USF1 expression is stabilized in the presence of USF2. **Panel A:** *USF2* KO Jurkat T cells were transduced with doxycycline (Dox) inducible Flag tagged USF2 expression vector. Following incubation with 1 μg/mL Dox for the indicated time, whole cell lysates were collected and analyzed by immunoblotting with antibodies against USF1, USF2, and Tubulin. **Panel B:** RNA was extracted from wildtype and *USF2* KO Jurkat cells and subject to Q-RT-PCR using primers specific for *USF2* mRNA. *USF2* transcript expression was normalized to *GAPDH* (*n* = 3, mean ±LSD). **Panel C:** Normalized RNA-seq *USF1* mRNA counts from WT and *USF2* KO Jurkat cells under basal (Ve) and T cell activated condition (PMA/Ion) (n = 3, mean ±LSD). **Panel D:** WT, *USF1* KO, or *USF2* KO cells were left untreated (Ve, DMSO) or stimulated by treatment with 5 nM PMA and 1 μM ionomycin. Following 4 hrs, whole lysates were collected and immunoblotted using antibodies against USF1, USF2, and Tubulin.

### USF2 stabilizes USF1 protein

Previously, we observed that depletion of USF2 in Jurkat T cells (Fig. 1*B*) (Fig. S1*C*) or HEK293T cells (Fig. 1*E*) invariably caused a corresponding decrease in USF1 protein abundance. To examine the mechanism by which USF1 protein abundance is dependent upon USF2, we first generated a Jurkat *USF2* KO cell line where Flag tagged USF2 is expressed from a doxycycline-inducible promoter. We found that expression of USF2-Flag in the *USF2* KO cell line, upon addition of doxycycline, rescued expression of USF1 protein to greater levels as compared to WT cells (Fig. 3*A*). We also observed that *USF2* KO did not cause a decrease in *USF1* mRNA abundance as compared to WT cells (Fig. 3*B*), regardless of T cell activation (Fig. 3*C*), indicating that differences in USF1 expression in *USF2* KO cells must relate to protein stability or effects on translation. Finally, PMA/ionomycin induced T cell activation did not alter USF1 or USF2 protein abundance in WT cells, and USF1 protein levels were decreased in *USF2* knockout cells regardless of cellular activation state (Fig. 3*D*).

### USF proteins can bind the HIV-1 LTR independently

We examined the requirements of USF1 and USF2 for binding to the HIV-1 LTR using chromatin immunoprecipitation and analysis by quantitative PCR (ChIP-qPCR). Because we were unable to immunoprecipitate USF1 with antibodies against native protein, we generated mHIV-Luc Jurkat T *USF1* and *USF2* KO cell lines that stably express Myc tagged USF1 (Fig. 4*A*). We found that USF1-Myc was associated with the LTR expressed in both the *USF1* and *USF2* KO cell lines (Fig. 4*B*). However, we did observe slightly higher amounts associated with all three sites on the LTR in the USF1 knockout line, which expresses endogenous USF2 (Fig. 4*B*). Similarly, we found that USF2 was associated with the LTR at equivalent amounts in WT and the *USF1* knockout lines (Fig. 4*C*). These results indicate that USF1 and USF2 are both capable of binding specific *cis*-elements on the LTR as homodimers *in vivo*, a result that is consistent with binding of recombinant USF1 and USF2 proteins *in vitro* (Malcolm et al. 2008) (Chen et al. 2005). Thus, although USF1 and USF2 were predominately shown to function as heterodimers *in vivo* (Sirito et al. 1994) (Sawadogo et al. 1988), USF2 is capable of binding independently of USF1 at multiple sites in the HIV-1 LTR for activation of transcription. In contrast, as indicated above, USF1 protein abundance is dependent upon USF2, and therefore is incapable of regulating HIV-1 transcription in *USF2* knockout cells.

**Figure 4.**
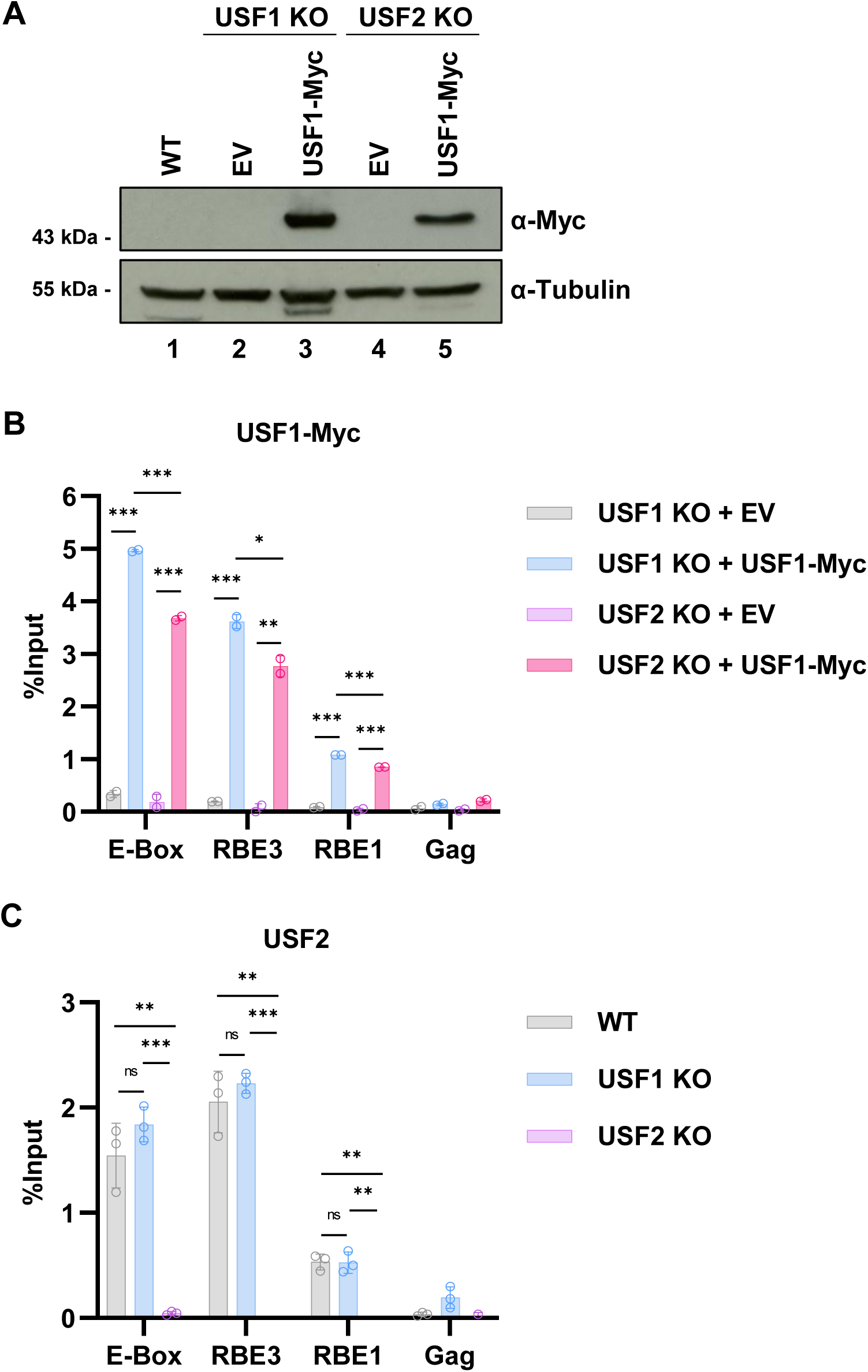
USF1 and USF2 are localize independently to the HIV-1 LTR. **Panel A:** *USF1* or *USF2* knockout cells were transduced with an empty vector (EV) or Myc tagged USF1 expression vector. Whole cell lysates were collected and immunoblotted with antibodies against Myc and Tubulin. **Panel B:** *USF1* or *USF2* knockout Jurkat T cells transduced with empty vector (EV) or USF1-Myc expression vector were subject to ChIP-qPCR using antibodies against Myc. ChIP-qPCR results are normalized by subtraction of values produced with sample paired non-specific IgG immunoprecipitation (*n* = 2, mean ±LSD). **Panel C:** WT or *USF1* or *USF2* knockouts were subject to ChIP-qPCR using antibodies targeting endogenous USF2. ChIP-qPCR results are normalized by subtraction of values produced with sample paired non-specific IgG immunoprecipitation (*n* = 2 - 3, mean ±LSD).

### Global regulation of transcription by USF1 and USF2 in unstimulated T cells

Given the surprising observation that Jurkat T cells with *USF1* and *USF2* gene knockouts have distinct effects on HIV-1 expression we sought to determine the degree that these factors cooperatively regulate global gene expression. To examine these effects, we performed RNA-seq on cell lines bearing *USF1* or *USF2* gene knockouts. In this analysis we identified 236 differentially expressed genes (DEG) upon *USF1* depletion and 563 DEG following *USF2* disruption (Fig. 5*A*, 5*B*). Of the genes affected by loss of *USF1*, slightly more were down-regulated than up- regulated (146 to 90, respectively), while *USF2* depletion primarily resulted in decreased expression of target genes (410 down-regulated to 153 up-regulated) (Fig. 5*A*, 5*B*). Comparison of transcriptomes from *USF1* and *USF2* KO cells demonstrated some overlap amongst down- regulated genes with 34% of *USF1* KO down-regulated genes similarly regulated in *USF2* KO cells (Fig. 5*C*). However, amongst upregulated genes we observed only 7% of genes upregulated upon USF1 depletion being likewise upregulated in *USF2* knockout cells (Fig. 5*D*). Overall, therefore the *USF1* and *USF2* knockouts cause a surprisingly small overlap in differential gene regulatory effects in unstimulated Jurkat T cells (Fig 5*E*). Furthermore, Gene Set Enrichment Analysis (GSEA) indicated only one pathway with less than 25% false discovery rate (FDR) shared between *USF1/2* KO, “TNFA signaling via NFκB”, which was negatively enriched (Fig. 5*F*, 5*G*). These results indicate that USF1 and USF2 share regulation for a minor subset of genes in unstimulated T cells.

**Figure 5.**
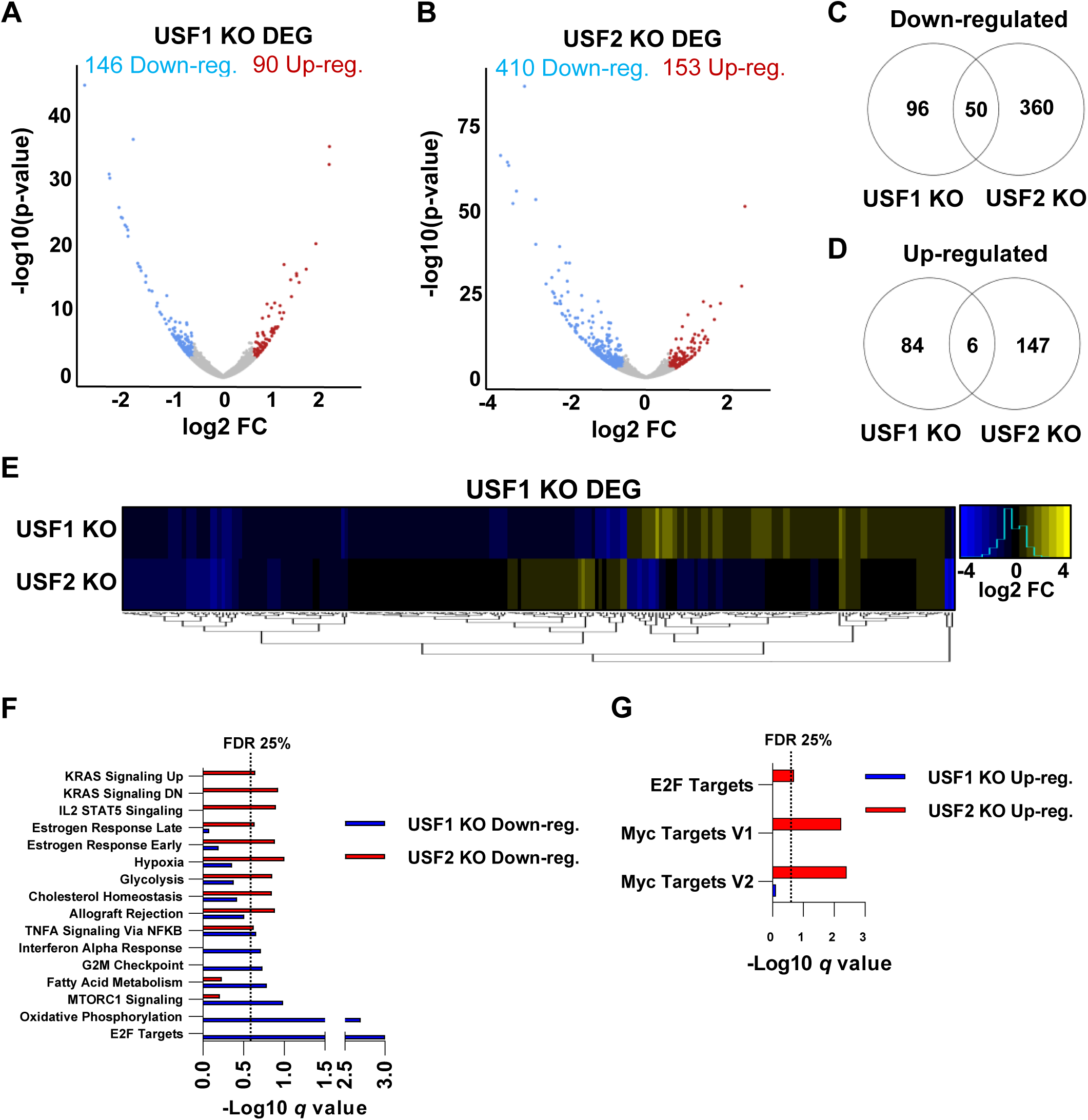
Transcriptomic effects of *USF1* or *USF2* knockout in unstimulated Jurkat T cells. **Panels A, B:** Volcano plots depicting DESeq2 analysis comparing WT and *USF1* KO (A) or *USF2* KO (B) Jurkat T cells. Analysis was performed on *n* = 3 RNA-seq samples with significant genes possessing p-value < 0.05 and fold change in expression > 1.5. **Panels C, D:** Venn diagram depiction of genes significantly down- (C) or up-regulated (D) upon *USF1* or *USF2* KO. **Panel E:** Heatmap depiction of the change in mRNA expression upon *USF1* or *USF2* KO of genes that were found to be differentially expressed upon *USF1* KO. **Panels F, G:** Gene set enrichment (GSEA) of Hallmark gene sets that are down-regulated (F) or up-regulated (G) for *USF1* or *USF2* KO transcriptomes.

### USF1 and USF2 cooperatively regulate global gene expression in activated T cells

USF1 and USF2 are signal regulated transcription factors (Galibert 2001) (Lupp et al. 2014) (Nowak et al. 2005) (Spohrer et al. 2017) that are phosphorylated upon T cell activation (Chen et al. 2005). Consequently, we also examined global gene expression in *USF1* and *USF2* knockout lines using RNA-seq from cells stimulated with PMA and ionomycin co-treatment. This analysis revealed a greater effect on global expression overall in stimulated cells compared to unstimulated, with 1846 and 2667 DEG upon loss of *USF1* and *USF2*, respectively (Fig. 6*A*, 6*B*). Importantly, depletion of either *USF1* or *USF2* resulted in the majority of DEG having decreased expression rather than increased (Fig. 6*A*, 6*B*). These results support the contention that USF1 and USF2 are T cell signal regulated factors that preferentially behave as transcriptional activators.

**Figure 6.**
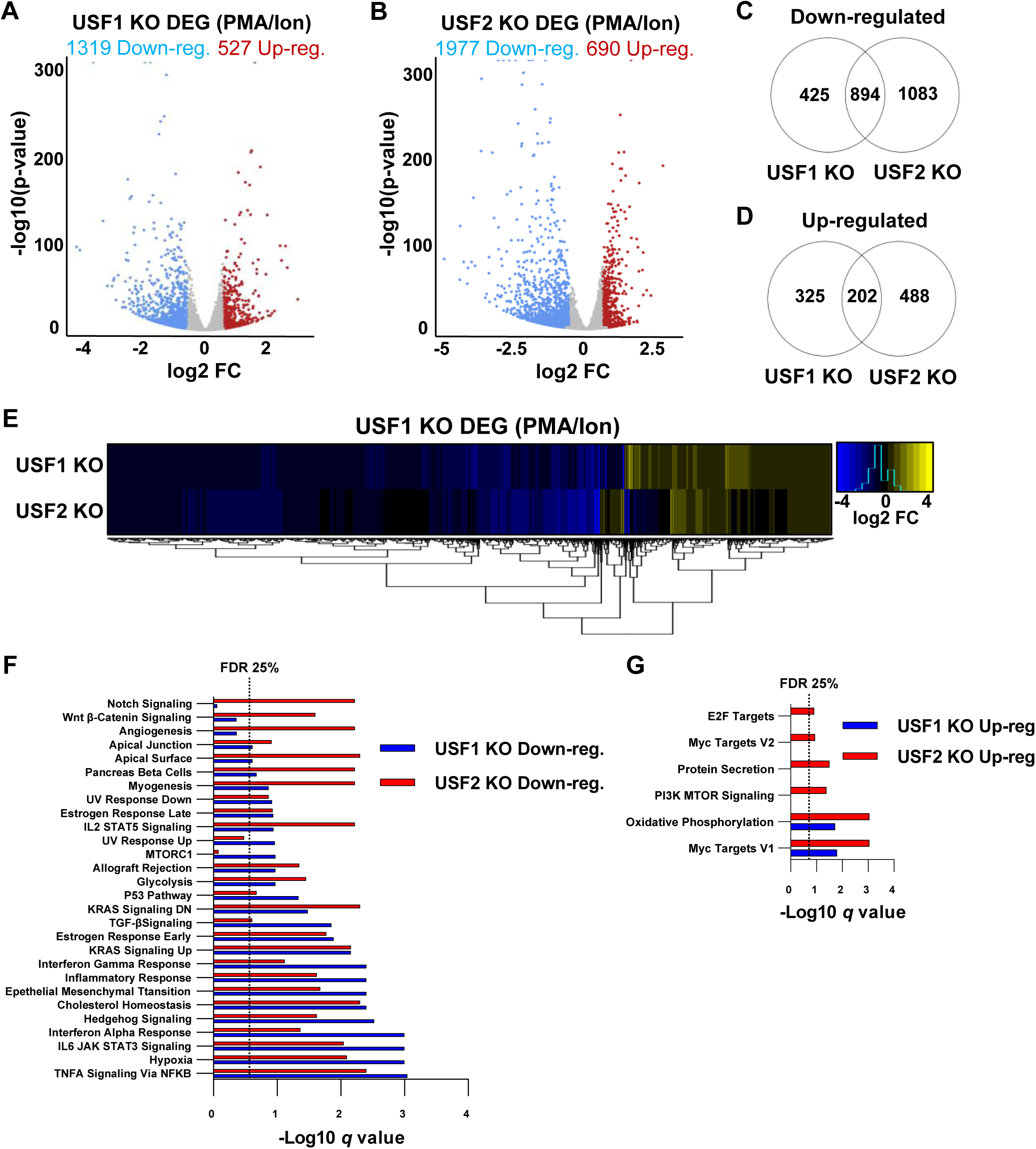
Effect of USF1 and USF2 knockout on the transcriptome of activated Jurkat T cells. **Panels A, B:** Volcano plots depicting DESeq2 analysis comparing WT and *USF1* KO (A) or *USF2* KO activated T cells (B). Jurkat T cells were stimulated by PMA/ionomycin co- treatment for 4 hrs prior to RNA extraction. Analysis was performed on *n* = 3 RNA-seq samples with significant genes possessing p-value < 0.05 and fold change in expression > 1.5. **Panels C, D:** Venn diagrams displaying genes significantly down- (C) or up-regulated (D) upon knockout of *USF1* or *USF2* under activating conditions. **Panel E:** Heatmap depiction of the change in gene expression upon *USF1* or *USF2* KO of genes that were differentially expressed upon *USF1* KO under activated T cell conditions. **Panels F, G:** Gene set enrichment (GSEA) of Hallmark gene sets that are down-regulated (F) or up-regulated (G) for *USF1*/ *USF2* KO transcriptomes following PMA/ionomycin mediated T cell activation.

Additionally, we observed a significantly greater proportion of genes that were similarly affected by *USF1* KO and *USF2* KO in activated T cells. In stimulated cells, we found that 68% of *USF1* KO downregulated and 38% of *USF1* KO upregulated genes were similarly mis- regulated upon *USF2* KO (Fig. 6*C*, 6*D*). Furthermore, a minority of USF1 DEG were found to be mis-regulated opposite that of *USF2* null cells (Fig. 6*E*). We performed GSEA analysis from *USF1* KO and *USF2* KO T cell transcriptomes to determine the predominant affected pathways, which revealed effects on genes involved in “TNFA Signaling *via* NFkB”, “Hypoxia”, and “Interferon Gamma Response” being negatively enriched (Fig. 6*F*), while “Oxidative Phosphorylation” and “Myc Targets V1” were found to be positively enriched in both *USF1* KO and *USF2* KO cell lines (Fig. 6*G*). Although genes from multiple pathways were similarly enriched in the *USF1* and *USF2* KO lines, we observed asymmetrical regulation between these factors as pathways including “mTORC1” and “Notch Signaling” were negatively enriched in *USF1* and *USF2* knockout cells, respectively (Fig. 6*F*). Interestingly, USF1 was previously implicated in regulating the UV and tanning response (Galibert 2001) (Corre et al. 2004) (Bouafia et al. 2014), yet we found that “UV Response Down” was negatively enriched in cells deficient of either USF1 or USF2 (Fig. 6*F*) suggesting that both factors mediate this process. Taken together, these results indicate that USF1 and USF2 display a high degree of cooperativity for global gene regulation during T cell activation. As noted previously, loss of USF2 results in a corresponding decrease in USF1 expression, and therefore it is possible that USF2 may partially compensate for loss of USF1 protein in both unstimulated and stimulated T cells.

### USF1 and USF2 facilitate T cell activation

Having observed that loss of either USF1 or USF2 generated an abundance of down-regulated genes upon T cell activation compared to wildtype, we examined the potential role of these factors in regulating inflammatory response. From RNA-seq analysis we identified 2790 down-regulated and 3481 up-regulated genes in response to T cell activation (Fig. 7*A*). Notably, genes that were induced in response to stimulation displayed a much greater range of expression compared to genes that were inhibited (Fig. 7*A*). We then examined the effect of *USF1* and *USF2* gene knockout on expression of genes that are activated or repressed upon T cell stimulation. Here we found that 755 (22%) and 1114 (32%) of genes induced upon T cell stimulation showed significantly less expression in the absence of USF1 and USF2 respectively (Fig. 7*B*). Genes that were repressed upon T cell activation were much less affected by *USF* disruption, with only 187 (7%) and 272 (10%) displaying greater expression upon loss of USF1 or USF2, respectively (Fig. 7*B*). These results are consistent with the above observations indicating the role of USF1 and USF2 as transcriptional activators.

**Figure 7.**
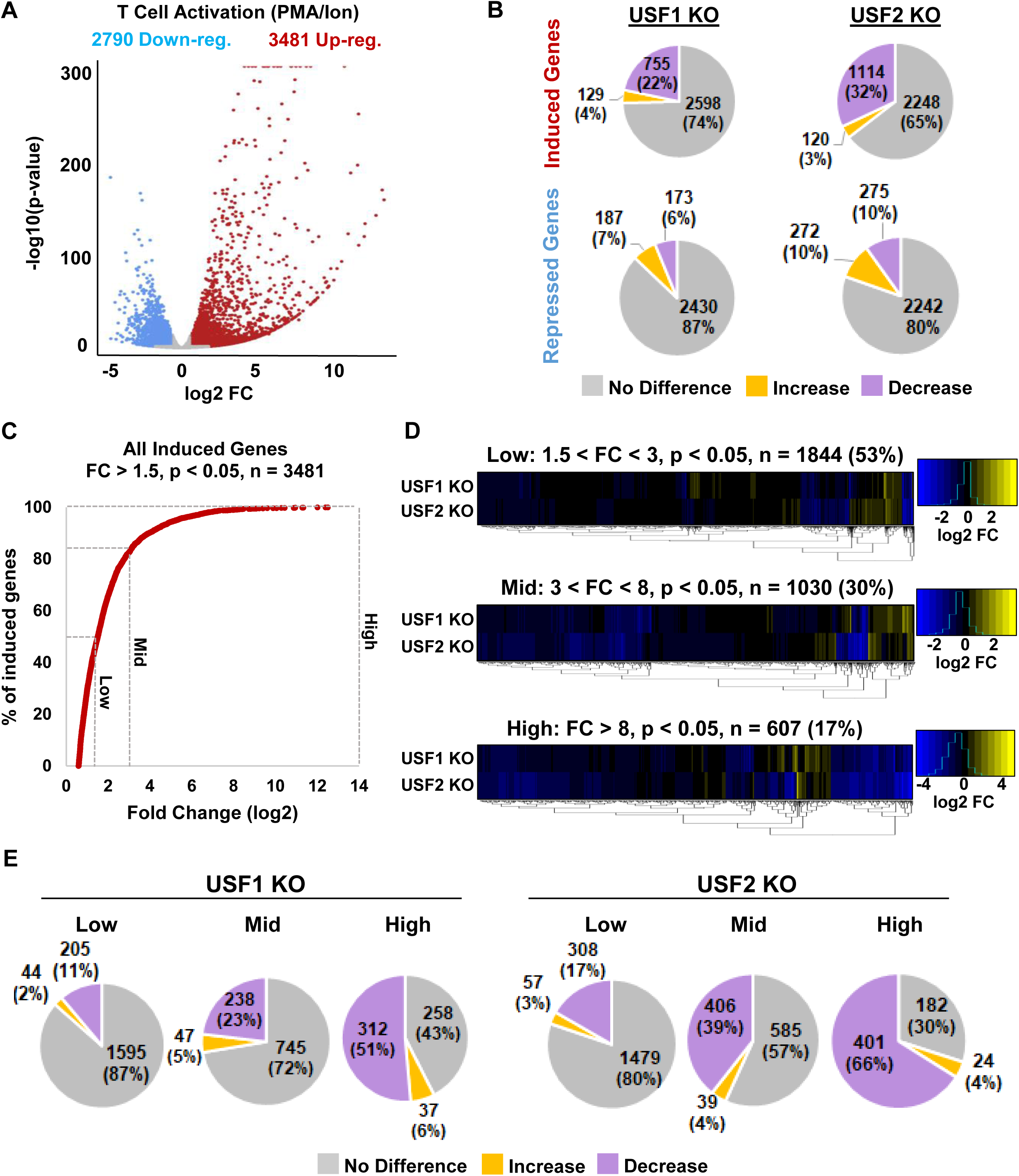
USF1 and USF2 facilitate T cell activation. **Panel A:** Volcano plot depiction of DESeq2 analysis comparing the transcriptomes of unstimulated and PMA/ionomycin activated Jurkat T cells. Analysis was performed on *n* = 3 RNA-seq samples with significant genes having p-value < 0.05 and fold change > 1.5. **Panel B:** Pie charts depicting the effect *USF1* (left) or *USF2* (right) knockout on T cell activation induced (top) or repressed (bottom) genes. **Panel C:** The number of T cell activation genes that are induced, shown as a percent of the total, given the fold change in expression. **Panel D:** Heatmap depicting the effects of *USF1,* and *USF2* KO on T cell activation induced genes. The T cell activation induced genes are stratified based on expression levels into Low, Mid, and High. **Panel E:** Pie charts depicting the effects of *USF1* or *USF2* KO on T cell activation genes that are induced at Low, Mid, and High expression levels.

T cell activation was previously shown to alter the expression of numerous genes with a wide degree of variance (Tong et al. 2016), consistent with our results indicating that nearly 50% of genes induced upon stimulation showing less than a 3-fold change in expression (Fig. 7C). Consequently, it is possible that genes displaying the large variance in induction are regulated by different mechanism(s). To examine this with respect to USF1 and USF2, we segmented genes induced upon T cell activation into three groups by level of induction, designated Low (fold change in expression between 1.5 and 3), Mid (fold change in expression between 3 and 8), and High (fold change greater than 8) (Fig. 7*C*). By stratifying genes based on level of induction, it became apparent that genes that were highly expressed in response to T cell stimulation were more potently inhibited in *USF1* or *USF2* KO cells (Fig. 7*D*). Furthermore, we found that more of the genes that were down regulated upon USF1 or USF2 depletion are those that are highly induced in response to T cell activation (Fig. 7*E*). Additionally, we found that *USF1* and *USF2* knockout cells displayed partially defective activation of genes downstream of key T cell signaling pathways including TNFα and Ras (Fig. S4). Collectively, these results indicate that USF1 and USF2 are signal regulated factors required for full induction of genes upon T cell activation, a finding that is consistent with the evolved capability of the HIV-1 LTR to pirate transcription factors downstream of T cell signaling (Sadowski and Mitchell 2005).

### USF2 deficient cells are refractory for HIV-1 infection

HIV-1 infection of resting CD4^+^ T cells is highly inefficient compared to infection of activated cells (Korin and Zack 1998) (Plesa et al. 2007) (Swiggard et al. 2005) (Unutmaz et al. 1999), and although Jurkat cells are actively dividing, they too are preferentially infected based on activation state (Matthew S Dahabieh et al. 2014). Because HIV-1 favors infection of stimulated T cells while loss of USF1 and USF2 dampens T cell activation, we sought to examine the role of these factors for HIV-1 infection. For this purpose, we created *USF1* and *USF2* knockouts in Jurkat T cells that did not contain a previously integrated provirus. Consistent with the results shown above, we observe loss of USF1 expression in both *USF1* and *USF2* knockout lines (Fig. 8*A*). These lines were infected with Red-Green-HIV-1 (RGH) (M. S. Dahabieh et al. 2013) or HIV_GKO_ (Battivelli and Verdin 2018) where GFP is expressed from the 5’ LTR and constitutive promoters express mCherry or mKO2 in RGH and HIV_GKO_, respectively (Fig. 8*B*). These dual reporter HIV-1 derivatives allow differentiation of productive and latent infections by flow cytometry (Fig. 8*C*, 8*D*) (Matthew S Dahabieh et al. 2014). We found that infection of *USF1* knockout cells produced a small but not significant decrease in RGH (Fig. 8*E*) or HIV_GKO_ (Fig. 8*F*) infectivity compared to WT, 4 days post-infection. In contrast, *USF2* knockout cells were infected roughly half that of wildtype cells by both reporter virus (Fig. 8*C*-8*F*). Interestingly, although USF2 is required for reactivation of latent provirus, we found no change in the proportion of cells that establish immediate latency upon infection of *USF2* KO T cells (Fig. 8*G*, 8*H*). Taken together, these observations indicate that USF2 is important for infection of T cells by HIV-1 by a process that likely involves cellular activation state.

**Figure 8.**
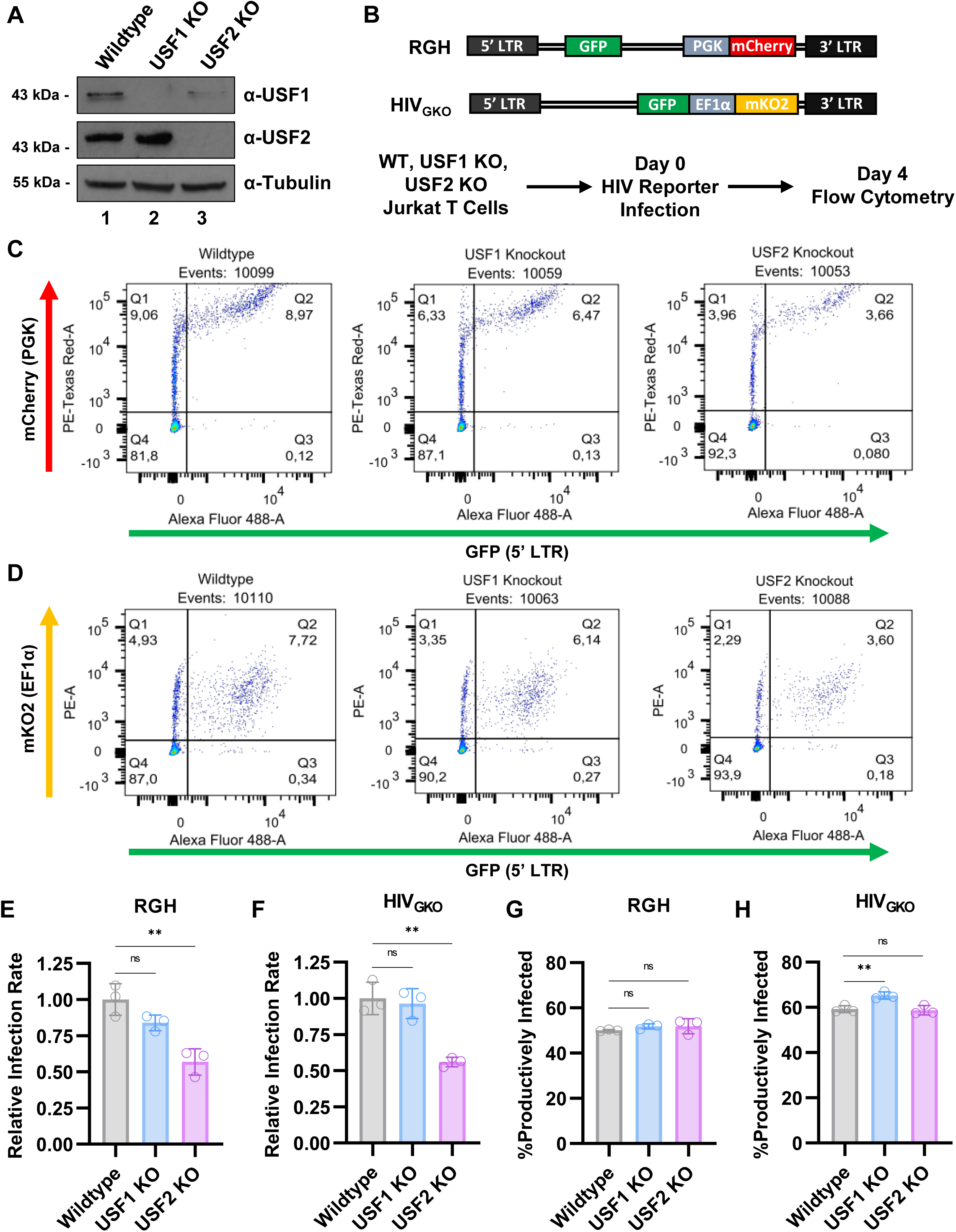
T cells deficient of USF2 are resistant to HIV-1 infection. **Panel A:** CRISPR-Cas9 was used to produce *USF1* or *USF2* knockout in Jurkat T cells. Whole cell lysates were subject to immunoblotting with antibodies against USF1, USF2, and Tubulin. **Panel B:** Schematic representation of the HIV-1 dual reporter viruses, and experimental summary. WT, *USF1* KO, or *USF2* KO Jurkat cells were infected with Red-Green-HIV-1 (RGH) or HIV_GKO_ dual-reporter viruses. Four days post- infection, proviral expression was determined by flow cytometry. **Panels C, D:** Representative FACS scatter plots of RGH (C) and HIV_GKO_ (D) infected cells, four days post-infection. Uninfected cells that are negative for fluorescence are in Q4, latent infections are located in Q1, productive infections are in Q2, and noise produced from viral recombination events are shown in Q3. **Panels E, F:** Relative infectivity was determined by flow cytometry four days post-infection of WT, *USF1* KO, or *USF2* KO Jurkat cells with equivalent amounts of RGH (E) or HIV_GKO_ (F) virions (*n* = 3, mean ±LSD). **Panels G, H:** As in E, F but the ratio of productive infections was analyzed by flow cytometry (*n* = 3, mean ±LSD).

### USF2 mediates establishment of HIV latency

We found that *USF2* knockout inhibits reactivation of latent HIV-1 provirus (Fig. 1) but does not affect the capacity of newly infected cells to immediately establish latent provirus (Fig. 8*G*, 8*H*). To further examine the role of USF for establishment of latency we monitored the proportion of productively infected cells for three weeks following infection. Previous observations indicate that the internal constitutively expressed mCherry and mKO2 reporters in cells infected with RGH and HIV_GKO_ become silenced following extended culture (M. S. Dahabieh et al. 2013) (Battivelli and Verdin 2018). Consequently, we isolated infected populations by fluorescent activated cell sorting (FACS) two days post infection (Fig. 9*A*), and then WT, *USF1* KO, and *USF2* KO infected cells were analyzed by flow cytometry over the following 22 days (Fig. 9*B*, 9*C*). Similar to the results with unsorted infected cells (Fig. 8), we found that WT, USF1, and USF2 deficient cells produced a nearly equivalent proportion of productively infected cells at 3- and 4-days post-infection (Fig. 9*D*, 9*E*). However, after 4-days infection, USF2 knockout cells accumulated latent HIV-1 infections at an accelerated rate compared to wildtype or *USF1* KO cells (Fig. 9*D*, 9*E*). These results indicate that USF2 promotes the expression of HIV-1 and encourages productive replication upon infection.

**Figure 9.**
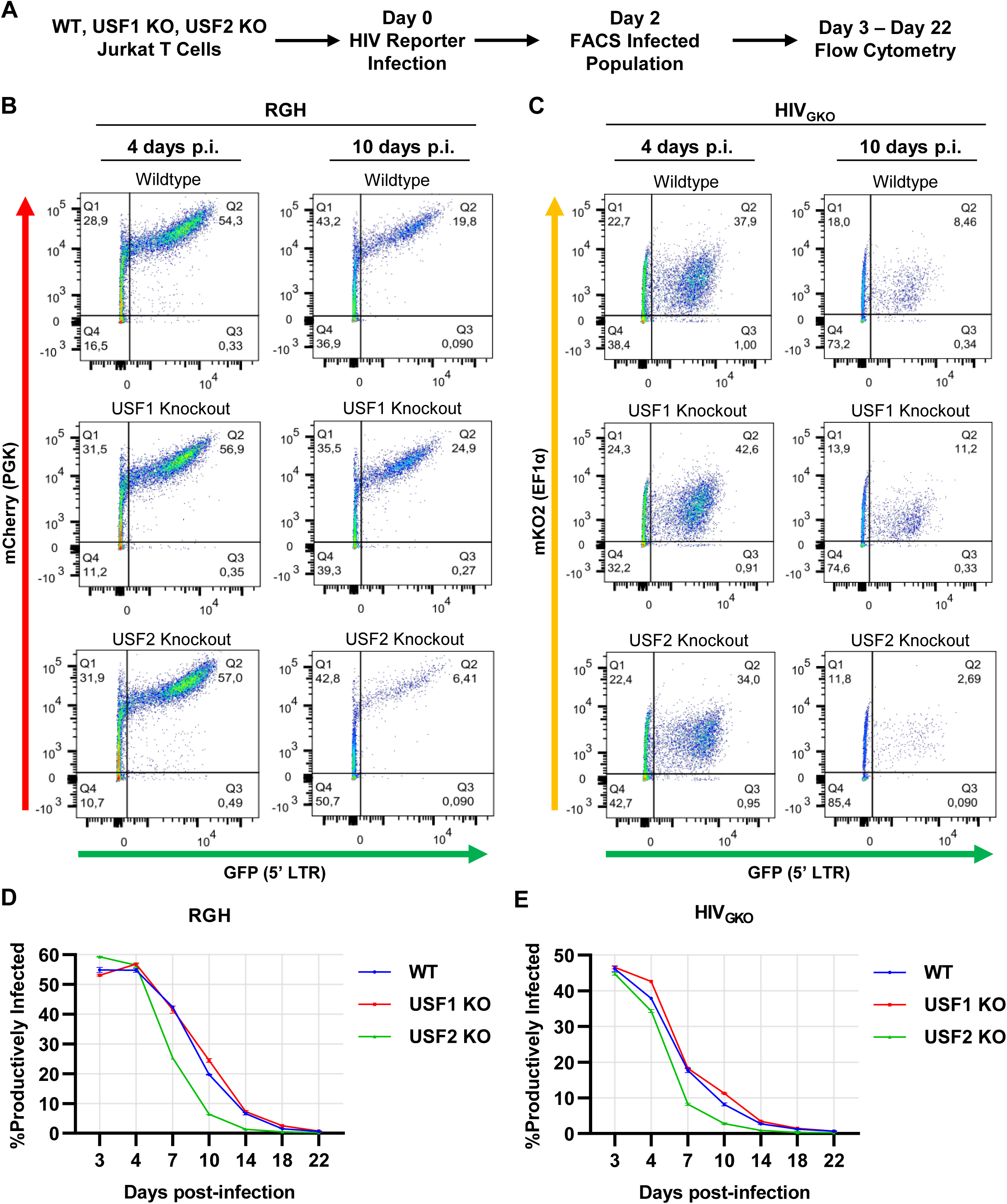
USF2 promotes the establishment of latency upon HIV-1 infection. **Panel A:** Schematic representation of the HIV-1 reporter assay. WT, *USF1* KO, or *USF2* KO Jurkat cells were infected with RGH or HIV_GKO_ dual-reporter virus at an M.O.I. of ∼0.1. Two days post- infection, latent and productively infected cells were isolated by FACS. Expression of HIV-1 reporter expression in the infected populations was examined over a 22-days by flow cytometry. **Panels B, C:** Representative flow cytometry scatter plots of RGH (B) or HIV_GKO_ (C) infected Jurkat cells at 4- and 10-days post-infection. **Panels D, E:** Summary of the percent of productive RGH (D) or HIV_GKO_ (E) infections as determined by flow cytometry at the indicated times post infection (*n* = 2, mean ±LSD).

## Discussion

The upstream stimulatory factor (USF) was among the first protein factors identified as a sequence specific DNA factor required for activation of transcription in eukaryotes (Sawadogo 1985) (Gregor, Sawadogo, and Roeder 1990), and among the first human factors discovered to bind the 5’ HIV-1 LTR (Du, Roy, and Roeder 1993). Various previous observations have indicated that USF1 and USF2 regulate transcription in response to multiple signaling pathways involving protein kinases, including at least MAPK/ ERKs (Corre et al. 2004) (Horbach et al. 2015), Casein kinase 2 (CK2) (Spohrer et al. 2017), Cdk2 (Cheung et al. 1999), protein kinase C (PKC) (Xiao, Kenessey, and Ojamaa 2002), DNA protein kinase (DNAPK) (Wong and Sul 2009), protein kinase A (PKA) (Sayasith, Lussier, and Sirois 2005), and Glycogen synthase kinase-3 (GSK-3) (Terragni et al. 2011). These protein kinases represent a significant fraction of signaling networks in human cells, which implicates USF1 and USF2 as key regulatory factors involved in coordinating gene expression in response to diverse cellular stimuli. USF-1 and USF-2 are ubiquitously expressed in human cells (Sirito et al. 1994), and consequently regulation of their function is likely cell type specific.

USF1/2 was found to bind the HIV-1 LTR at an E-box element in the upstream modulatory region at -160 to -165 from the transcriptional start site (Maekawa et al. 1991), where it was shown to activate transcription in cooperation with Ets-1 (Sieweke 1998). The upstream E-box site for USF is not particularly well conserved on LTRs from infected individuals who develop symptoms of AIDS (Mario Clemente Estable et al. 1996). USF1 and USF2 were subsequently found to form part of the factor RBF-2, which binds two highly conserved cis-elements at -160 (RBE3) and -20 (RBE1) (Bell and Sadowski 1996) (Matthew S. Dahabieh et al. 2011). These sites were initially identified because of their requirement for activation of HIV-1 transcription by activated RAS in T cells (Bell and Sadowski 1996); this factor was purified Jurkat nuclear extracts using oligonucleotide affinity (M. C. Estable et al. 1999), and found to contain USF1, USF2 and TFII-I (Chen et al. 2005). The RBE3 element is not typical of other characterized binding sites for USF1/2 (ACTGCTGA), and interaction of USF with this site *in vitro* and *in vivo* is dependent upon TFII-I (Malcolm et al. 2008) (Chen et al. 2005). In contrast RBE1 (CAGCTGC) (Bell and Sadowski 1996), near the transcriptional start, site resembles a typical E-box element (CANNTG), and consequently binding of USF1/2 to this site is less dependent upon TFII-I (Matthew S. Dahabieh et al. 2011). Consistent with our previous observations on RBF-2 we find that disruption of *USF2* in particular, inhibits expression of HIV-1 in response to T cell signaling as well as a variety of latency reversing agents (Fig. 1, 2). Furthermore, *USF2* disruption inhibits initial establishment of HIV-1 infections (Fig. 8) and reduces the proportion of infected cells that undergo productive gene expression (Fig. 9).

Mutation of RBE3 and/or RBE1 prevents induction of proviral expression in response to T cell signaling, and although USF1/2 and TFII-1 have been implicated in regulation of signal responsive gene expression (Horbach et al. 2015) (Segura-Puimedon et al. 2013), the mechanism(s) by which RBF-2 activates HIV-1 expression in response to T cell receptor engagement has not been elucidated. TFII-1, in association with serum response factor (SRF) was shown to regulate expression of the *c-fos* promoter in response to Ras-stimulated MAPK signaling by direct interaction with ERK1 (D.-W. Kim and Cochran 2000). Stimulation of T cells by antigen presentation causes activation of multiple pathways downstream of the T cell receptor, including the Ras-Raf-Mek-Erk (MAPK) pathway (Malcolm et al. 2007) (Sadowski and Mitchell 2005), and consequently the role of USF1/2 for activation of HIV-1 transcription likely includes, at least in part, interaction with TFII-I for binding of the conserved RBE1 and RBE3 sites on the LTR (Malcolm et al. 2008). Considering that USF1 and USF2 are reported to be regulated by multiple protein kinases as described above, it seems likely that these proteins mediate additional function(s) to regulate HIV-1 activation in response to T cell stimulation. Accordingly, we found that USF1, USF2 and TFII-I all become hyperphosphorylated in T cells stimulated with PMA and ionomycin (Chen et al. 2005).

From RNA-seq analysis, we found that *USF1* and *USF2* gene knockouts produce the most significant effect in cells stimulated with PMA and ionomycin, where we observe 1319 and 1977 downregulated genes, respectively (Fig. 6), whereas the knockouts have a significantly smaller effect in unstimulated cells (Fig. 5). This observation indicates that these factors play a significant role in regulating T cell activation response. Consistently, we observed that a significant proportion of genes that are highly induced in response to T cell activation were among those most seriously affected by disruption of USF1 and USF2 activation (Fig. 7*E*). Interestingly the most predominant classes of genes that were negatively impacted by *USF1* and *USF2* knockout in stimulated cells, as identified by gene set enrichment analysis included those involved in TNFα signaling through NFκB, signaling through Ras, and response to interferon γ (Fig. 6F) indicating an important role in regulation of inflammatory response. Important to note, because the USF factors seem to be ubiquitously expressed (Sirito et al. 1994), and their function may be modified by multiple protein kinases, the spectrum on

Despite that USF1 and USF2 are thought to predominately exist as heterodimers in most cell types (Sirito et al. 1998) (Sirito et al. 1994), we find that disruption of their genes cause significantly different alterations in the transcriptome, but where we observed greater overlap between differentially regulated genes in stimulated cells relative to unstimulated (Fig. 5, 6). Important to note, in stimulated cells we observed opposite effects on genes involved in mTORC1 and Notch signaling (Fig. 5F). These results are consistent with previous observations indicating distinct functions for USF1 and USF2. For example, *USF2* was associated with progression of leukemia through upregulation of *HOXA9* (Zhang et al. 2020). Additionally, overexpression of *USF2* was linked to rheumatoid arthritis that is refractory to treatment through the upregulation of proinflammatory cytokine expression (Hu et al. 2020). Phosphorylation of USF2 by Cdk5 was shown to stabilize this protein, which contributes to proliferation of prostate cancer cells (Chi et al. 2019). Additionally, differential binding of USF1 and USF2 to the plasminogen activator inhibitor type-1 promoter were reported to regulate expression during G_0_ to G_1_ cell cycle transition (Olave et al. 2010) (Qi et al. 2014). The N-terminal regions of USF-1 and USF-2 have ∼40% sequence identity (Sirito et al. 1998) (Sirito et al. 1994), and consequently because these factors are capable of binding the same E-box element (Sawadogo et al. 1988), alone or as heterodimers, the specific function of these factors has not been fully characterized. Our results indicate that *USF2* is required for USF1 protein expression and consequently this complicates previous analysis of differential effects of USF1 and USF2 for expression of specific genes. Considering this, it is remarkable that *USF1* and *USF2* gene knockouts produce significantly different transcriptome phenotypes.

We find that USF1 protein levels are significantly reduced upon *USF2* gene knockout or with shRNA silencing in Jurkat T cells and HEK293T cells. USF2 depletion does not affect *USF1* mRNA expression in either unstimulated or stimulated T cells, and consequently loss of USF2 must cause USF1 protein instability or inhibit translation of its mRNA. Overexpression of Flag tagged USF2 in *USF2* knockout cells restored USF1 protein levels, confirming that the presence of USF2 enables USF1 expression or stability. We propose that because USF1 and USF2 predominately exist as heterodimers (Sirito et al. 1994), this interaction stabilizes USF1 protein, and consequently loss of USF2 would result in USF1 protein instability. We have tried to examine this possibility, but this analysis was complicated by the finding that USF1 stability was not enhanced by the proteasome inhibitor MG132, or expression of the C-terminal region of USF2 containing the b-HLH leucine zipper motif. Nevertheless, such a mechanism regulating stability of heterodimer partners has been described previously, for the yeast MATa and α2 proteins are mutually stabilized by their heterodimer formation (Johnson et al. 1998).

Our results have revealed novel properties of the ubiquitously expressed USF2 protein, and a central role for regulation of HIV-1 gene expression. Aberrant expression of this factor was previously linked to additional human diseases, including rheumatoid arthritis (Hu et al. 2020), and cancer (Horbach et al. 2015). Consequently, a more detailed understanding of USF2 function and regulation will provide significant insight towards mechanisms regulating HIV-1 provirus expression. It will be particularly important to understand the role of USF2 for epigenetic regulation and chromatin organization at the HIV-1 LTR and cellular target genes, as well as the relative contribution of associated factors, including TFII-I for signal responsive gene expression.

## Materials and Methods

### Cell and virus culture

Jurkat E6-1, Jurkat Tat mHIV-Luciferase, and HEK293T cells were cultured under standard conditions of 37°C and 5% CO_2_ atmosphere. Jurkat cells were cultured in RPMI-1640 media while HEK293T were cultured in DMEM. Vesicular stomatitis virus G (VSV-G) pseudotyped viral stocks were produced by co-transfecting HEK293T cells with a combination of viral molecular clone, psPAX, and pVSV-G at a ratio of 8 μg : 4 μg : 2 μg. Transfections were performed with polyethylenimine (PEI) at a ratio of 6:1 (PEI:DNA) in Gibco Opti-MEM^TM^. Lentiviral infections were performed by plating 1x10^6^ cells in 24-well plates with media containing 8 μg/mL polybrene and the amount of viral supernatant to give the desired multiplicity of infection (M.O.I.) as indicated. Plates were subsequently spinoculated for 1.5 hrs at 1500 rpm.

### Immunoblotting

Western blotting was performed as previously described (Hashemi e*t al*, 2018). Antibodies were as follows: Tubulin (1:20000) – Abcam ab7291, Flag (1:20000) – Sigma Aldrich F3165, Myc (1:2500) – Santa Cruz Biotechnology sc-40, USF1 (1:500) – Santa Cruz Biotechnology sc-390027, USF2 (1:8000) – Abcam ab264330, H3 (1:2000) - Abcam ab24834, Goat Anti-Rabbit-HRP – Abcam ab6721 (1:2000000), Goat Anti-Mouse-HRP – Pierce 1858413 (1:20000).

### Luciferase expression assays

For Jurkat Tat mHIV-Luciferase expression assays, 1x10^5^ of the luciferase expressing cells were plated with 100 µL media in 96-well plates. Luciferase activity was measured after the indicated time of treatment. Measurements were performed using Superlight^TM^ luciferase reporter Gene Assay Kit (BioAssay Systems) as per the manufacturer’s instructions; 96 well plates were read in a VictorTM X3 Multilabel Plate Reader.

### Transfection assays

6.67x10^5^ Wildtype or *USF2* KO HEK293T cells were plated in 2 mL DMEM in 6-well plates. The following day, cells were transfected with 0.5 ng of the indicated LAI LTR-GFP reporter construct and 1 μg of the indicated EF1α-RFP construct. Transfections were performed with polyethylenimine (PEI) at a ratio of 1:4 (DNA:PEI) in Gibco Opti-MEM^tm^. One day post-transfection, cells were detached with trypsin-EDTA and analyzed by flow cytometry.

### Flow cytometry

Cells were treated as indicated in the figure legends. Following the indicated treatment, Jurkat derived cells were suspended in PBS while HEK293T derived cells were suspended in PBS containing 10% trypsin-EDTA to prevent aggregation. Flow cytometric analysis was performed on a BD Biosciences LSRII-561 system where threshold forward scatter (FSC) and side scatter (SSC) parameters were set so that a homogenous population of live cells was counted (Fig. S5). FlowJo software (TreeStar) was used to analyze data and determine the indicated mean fluorescent intensity (MFI).

### Generation of USF1 and USF2 knockout clonal lines

*USF1* KO and *USF2* KO Jurkat Tat mHIV-Luciferase and wildtype Jurkat E6-1 clonal cell lines were generated by CRISPR-Cas9 gene editing. 2x10^6^ cells were co-transfected with Cas9 and gRNA targeting expression vectors specific for genomic *USF1* or *USF2.* Transfection was performed using the Neon Transfection System (Invitrogen) as per the manufacturer’s instructions with the following settings: Voltage, 1350 V; Width, 20 ms; Pulse number, 3x. For *USF1* KO, Jurkat cells were co-transfected with Cas9-BFP (pU6-CBh-Cas9-T2A-BFP: Addgene 64323) and gRNA (pSPgRNA: Addgene 47108) containing gRNA that targeted GCCAGGTAAGGGAGGGGGCC and GGAAGACGTACTTGACGTTG. For, *USF2* KO, Jurkat cells were co-transfected with Cas9-mCherry (pU6-CBh-Cas9-T2A-mCherry: Addgene 64324) and gRNA-BFP (pKLV2.2-h7SKgRNA-hU6gRNA-PGKpuroBFP: Addgene 72666) containing gRNA that targeted GTCGTGGCTGCCAGGGGCAC, GGAGGAGGGCGTCGAGCTGC, and GGCGGCCGAGGCTGTCAGCG. For *USF1* KO, transfected cells were isolated by live sorting (Astrios Flow Cytometer) BFP+ cells into 96-well plates containing complete RPMI-1640 while for *USF2* KO, mCherry+/BFP+ cells were isolated. Clones were expanded, and knockout of *USF1* and *USF2* was validated by PCR genotyping and western blotting.

To generate *USF2* KO HEK293T clonal cell lines, cells were co-transfected using polyethylenimine (PEI) with Cas9 (pU6-CBh-Cas9-T2A-BFP: Addgene 64323) and gRNA-Puro (pKLV2.2-h7SKgRNA-hU6gRNA-PGKpuroBFP: Addgene 72666) containing gRNA that targeted GGAGGAGGGCGTCGAGCTGC and GGCGGCCGAGGCTGTCAGCG. Briefly, 6.67x10^5^ HEK293T cells were plated with 2 mL DMEM in a 6-well plate. The following day, transfection was performed with 3 μg Cas9, 1 μg gRNA-Puro, and 12 μg PEI. Transfected cells were selected with 1 μg/mL puromycin for four days at which point single cells were plated in wells of 96-well plates by limiting dilution. Clones were expanded and *USF2* KO was validated by western blotting.

### USF1 and USF2 shRNA knockdown

Jurkat mHIV-Luciferase cells were infected with pLKO empty vector or pLKO shRNA expressing lentivirus at a M.O.I. ∼10. shRNA transduced cells were cultured for up to 8 days with 3 µg/mL puromycin, during which pools of puromycin selected cells were prepared for the indicated analysis. MISSION shRNA clones (Sigma) used for knockdown are as follows: USF1, TRCN0000020679 – GCTGGATACTGGACACACTAA (3’ UTR); USF2-1, TRCN0000020734 – TCCTCCACTTGGAAACGGTAT (3’ UTR); USF2-2, TRCN0000020737 – TCCAGACTGTAACGCAGACAA (CDS).

### ChIP-qPCR

ChIP-qPCR was performed as in Horvath *et al*, 2023 (Horvath et al. 2023) and Horvath *et al*, 2023 (Horvath, Brumme, and Sadowski 2023). Antibodies used are as follows: Myc (10 μg) – Santa Cruz Biotechnology sc-40, USF2 (10 μg) – Abcam ab264330.

### RT-PCR

RNA was extracted from Jurkat Tat mHIV-Luciferase cells following the indicated treatment using RNeasy Kit (Qiagen). RNA was analyzed using the Quant Studio 3 Real-Time PCR system (Applied Biosystems) using *Power* SYBR® Green RNA-to-CT™ 1-Step Kit (Thermo Fisher) as per the manufacturer’s instructions. RT-PCR data was normalized to *GAPDH* expression using the ΔΔCt method. Primers used are as follows: USF1 Fwd 5’ GCTCTATGGAGAGCACCAAGTC, Rev 5’ AGACAAGCGGTGGTTACTCTGC; GAPDH Fwd 5’ TGCACCACCAACTGCTTAGC, Rev 5’ GGCATGGACTGTGGTCATGAG.

### RNA-Seq

RNA was extracted from wildtype, *USF1* KO, or *USF2* KO Jurkat Tat mHIV-Luciferase human T cells that were left untreated or stimulated with PMA/ ionomycin for 4 hrs. Sample quality control was performed using an Agilent 2100 Bioanalyzer. Qualifying samples were prepped using NEBnext Ultra ii Stranded mRNA (New England Biolabs) standard protocol. Sequencing was performed on the Illumina NextSeq 500 with Paired End 42bp × 42bp reads. De-multiplexed read sequences were uploaded to the Galaxy web platform (Afgan e*t al*, 2018) and aligned to the hg38 reference genome using STAR. Subsequently, transcript assembly was accomplished with featureCounts and differential gene expression (DEG) was determined by DESeq2 analysis. DEG are defined as having a fold change in expression > 1.5 and p-value < 0.05 as to the compared condition. Volcano plots were generated on the Galaxy web platform and gene expression heatmaps were created in RStudio.

### Statistics and reproducibility

All replicates are independent biological samples and results are presented as mean values withL± standard deviation shown by error bars. The number of times that an experiment was performed is indicated in the figure legends. P-values were determined by performing unpaired samples *t*-test with the use of GraphPad Prism 9.0.0. Statistical significance is indicated at \**P* < 0.05, \*\**P* < 0.01, or \*\*\**P* < 0.001, with n.s. denoting non-significant *P* ≥ 0.05.

## Data availability

All data supporting the findings of this study are available within the article or from the corresponding author upon reasonable request (I. Sadowski). All high-throughput RNA-seq data has been deposited with NCBI GEO, accession GSE227850.

## Acknowledgments

We thank Andy Johnson and Justin Wong of the UBC Flow Cytometry Facility for performing FACS analysis as well as for assistance with flow cytometry. This research was supported by program project grant F16-01210, from the Canadian Institutes of Health Research.

## Author Contributions

Horvath R. M. performed all experiments. Experimental design was conceived by Horvath R. M. and Sadowski I. Horvath R. M. and Sadowski I. wrote the manuscript.

## References

1. Abner, Erik, and Albert Jordan. 2019. “HIV ‘Shock and Kill’ Therapy: In Need of Revision.” Antiviral Research 166 (June): 19–34. https://doi.org/10.1016/j.antiviral.2019.03.008.

2. Andrews, G. K. 2001. “The Transcription Factors MTF-1 and USF1 Cooperate to Regulate Mouse Metallothionein-I Expression in Response to the Essential Metal Zinc in Visceral Endoderm Cells during Early Development.” The EMBO Journal 20 (5): 1114–22. https://doi.org/10.1093/emboj/20.5.1114.

3. Battivelli, Emilie, and Eric Verdin. 2018. “HIVGKO: A Tool to Assess HIV-1 Latency Reversal Agents in Human Primary CD4+ T Cells.” BIO-PROTOCOL 8 (20). https://doi.org/10.21769/BioProtoc.3050.

4. Baxevanis, Andreas D., and Charles R. Vinson. 1993. “Interactions of Coiled Coils in Transcription Factors: Where Is the Specificity?” Current Opinion in Genetics & Development 3 (2): 278–85. https://doi.org/10.1016/0959-437X(93)90035-N.

5. Bell, B., and I. Sadowski. 1996. “Ras-Responsiveness of the HIV-1 LTR Requires RBF-1 and RBF-2 Binding Sites.” Oncogene 13 (12): 2687–97.

6. Bernhard, Wendy, Kris Barreto, Sheetal Raithatha, and Ivan Sadowski. 2013. “An Upstream YY1 Binding Site on the HIV-1 LTR Contributes to Latent Infection.” Edited by Peter Sommer. PLoS ONE 8(10): e77052. https://doi.org/10.1371/journal.pone.0077052.

7. Bernhard, Wendy, Kris Barreto, Amy Saunders, Matthew S. Dahabieh, Pauline Johnson, and Ivan Sadowski. 2011. “The Suv39H1 Methyltransferase Inhibitor Chaetocin Causes Induction of Integrated HIV-1 without Producing a T Cell Response.” FEBS Letters 585 (22): 3549–54. https://doi.org/10.1016/j.febslet.2011.10.018.

8. Bouafia, Amine, Sébastien Corre, David Gilot, Nicolas Mouchet, Sharon Prince, and Marie-Dominique Galibert. 2014. “P53 Requires the Stress Sensor USF1 to Direct Appropriate Cell Fate Decision.” Edited by Marshall S. Horwitz. PLoS Genetics 10(5): e1004309. https://doi.org/10.1371/journal.pgen.1004309.

9. Brooks, D. G., P. A. Arlen, L. Gao, C. M. R. Kitchen, and J. A. Zack. 2003. “Identification of T Cell-Signaling Pathways That Stimulate Latent HIV in Primary Cells.” Proceedings of the National Academy of Sciences 100 (22): 12955–60. https://doi.org/10.1073/pnas.2233345100.

10. Chen, J., T. Malcolm, M. C. Estable, R. G. Roeder, and I. Sadowski. 2005. “TFII-I Regulates Induction of Chromosomally Integrated Human Immunodeficiency Virus Type 1 Long Terminal Repeat in Cooperation with USF.” Journal of Virology 79 (7): 4396–4406. https://doi.org/10.1128/JVI.79.7.4396-4406.2005.

11. Cheung, Edwin, Petra Mayr, Federico Coda-Zabetta, Phillip G Woodman, and David S W Boam. 1999. “DNA-Binding Activity of the Transcription Factor Upstream Stimulatory Factor 1 (USF- 1) Is Regulated by Cyclin-Dependent Phosphorylation,” 8.

12. Chi, Tabughang, Tina Horbach, Claudia Götz, Thomas Kietzmann, and Elitsa Dimova. 2019. “Cyclin-Dependent Kinase 5 (CDK5)-Mediated Phosphorylation of Upstream Stimulatory Factor 2 (USF2) Contributes to Carcinogenesis.” Cancers 11 (4): 523. https://doi.org/10.3390/cancers11040523.

13. Corre, Sébastien, Aline Primot, Elena Sviderskaya, Dorothy C. Bennett, Sophie Vaulont, Colin R. Goding, and Marie-Dominique Galibert. 2004. “UV-Induced Expression of Key Component of the Tanning Process, the POMC and MC1R Genes, Is Dependent on the p-38-Activated Upstream Stimulating Factor-1 (USF-1).” Journal of Biological Chemistry 279 (49): 51226–33. https://doi.org/10.1074/jbc.M409768200.

14. Dahabieh, M. S., M. Ooms, V. Simon, and I. Sadowski. 2013. “A Doubly Fluorescent HIV-1 Reporter Shows That the Majority of Integrated HIV-1 Is Latent Shortly after Infection.” Journal of Virology 87 (8): 4716–27. https://doi.org/10.1128/JVI.03478-12.

15. Dahabieh, Matthew S, Marcel Ooms, Chanson Brumme, Jeremy Taylor, P Richard Harrigan, Viviana Simon, and Ivan Sadowski. 2014. “Direct Non-Productive HIV-1 Infection in a T-Cell Line Is Driven by Cellular Activation State and NFκB.” Retrovirology 11 (1): 17. https://doi.org/10.1186/1742-4690-11-17.

16. Dahabieh, Matthew S., Marcel Ooms, Tom Malcolm, Viviana Simon, and Ivan Sadowski. 2011. “Identification and Functional Analysis of a Second RBF-2 Binding Site within the HIV-1 Promoter.” Virology 418 (1): 57–66. https://doi.org/10.1016/j.virol.2011.07.002.

17. Desimio, Maria Giovanna, Erica Giuliani, and Margherita Doria. 2017. “The Histone Deacetylase Inhibitor SAHA Simultaneously Reactivates HIV-1 from Latency and up-Regulates NKG2D Ligands Sensitizing for Natural Killer Cell Cytotoxicity.” Virology 510 (October): 9–21. https://doi.org/10.1016/j.virol.2017.06.033.

18. Du, H., A.L. Roy, and R.G. Roeder. 1993. “Human Transcription Factor USF Stimulates Transcription through the Initiator Elements of the HIV-1 and the Ad-ML Promoters.” The EMBO Journal 12 (2): 501–11. https://doi.org/10.1002/j.1460-2075.1993.tb05682.x.

19. Estable, M. C., M. Hirst, B. Bell, M. V. O’Shaughnessy, and I. Sadowski. 1999. “Purification of RBF-2, a Transcription Factor with Specificity for the Most Conserved Cis-Element of Naturally Occurring HIV-1 LTRs.” Journal of Biomedical Science 6 (5): 320–32. https://doi.org/10.1007/bf02253521.

20. Estable, Mario Clemente, Brendan Bell, Abderrazzak Merzouki, Julio S G Montaner, Michael V O’Shaughnessy, and Ivan J Sadowski. 1996. “Human Immunodeficiency Virus Type 1 Long Terminal Repeat Variants from 42 Patients Representing All Stages of Infection Display a Wide Range of Sequence Polymorphism and Transcription Activity.” J. VIROL. 70: 10.

21. Fagagna, F d’Adda di, G Marzio, M I Gutierrez, L Y Kang, A Falaschi, and M Giacca. 1995. “Molecular and Functional Interactions of Transcription Factor USF with the Long Terminal Repeat of Human Immunodeficiency Virus Type 1.” Journal of Virology 69 (5): 2765–75. https://doi.org/10.1128/jvi.69.5.2765-2775.1995.

22. Finzi, D. 1997. “Identification of a Reservoir for HIV-1 in Patients on Highly Active Antiretroviral Therapy.” Science 278 (5341): 1295–1300. https://doi.org/10.1126/science.278.5341.1295.

23. Galibert, M.-D. 2001. “The Usf-1 Transcription Factor Is a Novel Target for the Stress- Responsive P38 Kinase and Mediates UV-Induced Tyrosinase Expression.” The EMBO Journal 20 (17): 5022–31. https://doi.org/10.1093/emboj/20.17.5022.

24. Gregor, P D, M Sawadogo, and R G Roeder. 1990. “The Adenovirus Major Late Transcription Factor USF Is a Member of the Helix-Loop-Helix Group of Regulatory Proteins and Binds to DNA as a Dimer.” Genes & Development 4 (10): 1730–40. https://doi.org/10.1101/gad.4.10.1730.

25. Groenen, Peter M.A., Emilio Garcia, Philippe Debeer, Koen Devriendt, Jean Pierre Fryns, and Wim J.M.Van de Ven. 1996. “Structure, Sequence, and Chromosome 19 Localization of HumanUSF2and Its Rearrangement in a Patient with Multicystic Renal Dysplasia.” Genomics 38 (2): 141–48. https://doi.org/10.1006/geno.1996.0609.

26. Horbach, Tina, Claudia GÃ¶tz, Thomas Kietzmann, and Elitsa Y. Dimova. 2015. “Protein Kinases as Switches for the Function of Upstream Stimulatory Factors: Implications for Tissue Injury and Cancer.” Frontiers in Pharmacology 6 (February). https://doi.org/10.3389/fphar.2015.00003.

27. Horvath, Riley M., Zabrina L. Brumme, and Ivan Sadowski. 2023. “Inhibition of the TRIM24 Bromodomain Reactivates Latent HIV-1.” Scientific Reports 13 (1): 556. https://doi.org/10.1038/s41598-023-27765-3.

28. Horvath, Riley M., Matthew Dahabieh, Tom Malcolm, and Ivan Sadowski. 2023. “TRIM24 Controls Induction of Latent HIV-1 by Stimulating Transcriptional Elongation.” Communications Biology 6 (1): 86. https://doi.org/10.1038/s42003-023-04484-z.

29. Hu, Dan, Emily C. Tjon, Karin M. Andersson, Gabriela M. Molica, Minh C. Pham, Brian Healy, Gopal Murugaiyan, et al. 2020. “Aberrant Expression of USF2 in Refractory Rheumatoid Arthritis and Its Regulation of Proinflammatory Cytokines in Th17 Cells.” Proceedings of the National Academy of Sciences 117 (48): 30639–48. https://doi.org/10.1073/pnas.2007935117.

30. Johnson, P. R., R. Swanson, L. Rakhilina, and M. Hochstrasser. 1998. “Degradation Signal Masking by Heterodimerization of MATalpha2 and MATa1 Blocks Their Mutual Destruction by the Ubiquitin-Proteasome Pathway.” Cell 94 (2): 217–27. https://doi.org/10.1016/s0092-8674(00)81421-x.

31. Joos, B., M. Fischer, H. Kuster, S. K. Pillai, J. K. Wong, J. Boni, B. Hirschel, et al. 2008. “HIV Rebounds from Latently Infected Cells, Rather than from Continuing Low-Level Replication.” Proceedings of the National Academy of Sciences 105 (43): 16725–30. https://doi.org/10.1073/pnas.0804192105.

32. Kedei, Noemi, Daniel J. Lundberg, Attila Toth, Peter Welburn, Susan H. Garfield, and Peter M. Blumberg. 2004. “Characterization of the Interaction of Ingenol 3-Angelate with Protein Kinase C.” Cancer Research 64 (9): 3243–55. https://doi.org/10.1158/0008-5472.CAN-03-3403.

33. Kim, D.-W., and B. H. Cochran. 2000. “Extracellular Signal-Regulated Kinase Binds to TFII-I and Regulates Its Activation of the c-Fos Promoter.” Molecular and Cellular Biology 20 (4): 1140–48. https://doi.org/10.1128/MCB.20.4.1140-1148.2000.

34. Kim, Youry, Jenny L. Anderson, and Sharon R. Lewin. 2018. “Getting the ‘Kill’ into ‘Shock and Kill’: Strategies to Eliminate Latent HIV.” Cell Host & Microbe 23 (1): 14–26. https://doi.org/10.1016/j.chom.2017.12.004.

35. Korin, Yael D., and Jerome A. Zack. 1998. “Progression to the G _1_ b Phase of the Cell Cycle Is Required for Completion of Human Immunodeficiency Virus Type 1 Reverse Transcription in T Cells.” Journal of Virology 72 (4): 3161–68. https://doi.org/10.1128/JVI.72.4.3161-3168.1998.

36. Li, Zichong, Jia Guo, Yuntao Wu, and Qiang Zhou. 2013. “The BET Bromodomain Inhibitor JQ1 Activates HIV Latency through Antagonizing Brd4 Inhibition of Tat-Transactivation.” Nucleic Acids Research 41 (1): 277–87. https://doi.org/10.1093/nar/gks976.

37. Lupp, Sarah, Claudia Götz, Sunia Khadouma, Tina Horbach, Elitsa Y. Dimova, Anna-Maria Bohrer, Thomas Kietzmann, and Mathias Montenarh. 2014. “The Upstream Stimulatory Factor USF1 Is Regulated by Protein Kinase CK2 Phosphorylation.” Cellular Signalling 26 (12): 2809–17. https://doi.org/10.1016/j.cellsig.2014.08.028.

38. Maekawa, Toshio, Tatsuhiko Sudo, Masashi Kurimoto, and Shunsuke Ishii. 1991. “USF-Related Transcription Factor, HIV-TF1, Stimulates Transcription of Human Immunodeficiency Virus-1.” Nucleic Acids Research 19 (17): 4689–94. https://doi.org/10.1093/nar/19.17.4689.

39. Malcolm, Tom, Jiguo Chen, Carol Chang, and Ivan Sadowski. 2007. “Induction of Chromosomally Integrated HIV-1 LTR Requires RBF-2 (USF/TFII-I) and RAS/MAPK Signaling.” Virus Genes 35 (2): 215–23. https://doi.org/10.1007/s11262-007-0109-9.

40. Malcolm, Tom, Joanna Kam, Pouya Sadeghi Pour, and Ivan Sadowski. 2008. “Specific Interaction of TFII-I with an Upstream Element on the HIV-1 LTR Regulates Induction of Latent Provirus.” FEBS Letters 582 (28): 3903–8. https://doi.org/10.1016/j.febslet.2008.10.032.

41. Nowak, Maxime, Audrey Helleboid-Chapman, Heidelinde Jakel, Geneviève Martin, Daniel Duran-Sandoval, Bart Staels, Edward M. Rubin, et al. 2005. “Insulin-Mediated Down- Regulation of Apolipoprotein A5 Gene Expression through the Phosphatidylinositol 3-Kinase Pathway: Role of Upstream Stimulatory Factor.” Molecular and Cellular Biology 25 (4): 1537–48. https://doi.org/10.1128/MCB.25.4.1537-1548.2005.

42. Olave, Nélida C., Maximiliano H. Grenett, Martin Cadeiras, Hernan E. Grenett, and Paul J. Higgins. 2010. “Upstream Stimulatory Factor-2 Mediates Quercetin-Induced Suppression of PAI-1 Gene Expression in Human Endothelial Cells.” Journal of Cellular Biochemistry 111 (3): 720–26. https://doi.org/10.1002/jcb.22760.

43. Pasquereau, Sébastien, and Georges Herbein. 2022. “CounterAKTing HIV: Toward a ‘Block and Clear’ Strategy?” Frontiers in Cellular and Infection Microbiology 12 (February): 827717. https://doi.org/10.3389/fcimb.2022.827717.

44. Pereira, L. A. 2000. “SURVEY AND SUMMARY A Compilation of Cellular Transcription Factor Interactions with the HIV-1 LTR Promoter.” Nucleic Acids Research 28 (3): 663–68. https://doi.org/10.1093/nar/28.3.663.

45. Petravic, Janka, Thomas A. Rasmussen, Sharon R. Lewin, Stephen J. Kent, and Miles P. Davenport. 2017. “Relationship between Measures of HIV Reactivation and Decline of the Latent Reservoir under Latency-Reversing Agents.” Edited by Guido Silvestri. Journal of Virology 91 (9): e02092–16, /jvi/91/9/e02092-16.atom. https://doi.org/10.1128/JVI.02092-16.

46. Plesa, Gabriela, Jihong Dai, Cliff Baytop, James L. Riley, Carl H. June, and Una O’Doherty. 2007. “Addition of Deoxynucleosides Enhances Human Immunodeficiency Virus Type 1 Integration and 2LTR Formation in Resting CD4 ^+^ T Cells.” Journal of Virology 81 (24): 13938–42. https://doi.org/10.1128/JVI.01745-07.

47. Puertas, Maria C, George Ploumidis, Michalis Ploumidis, Emilio Fumero, Bonaventura Clotet, Charles M Walworth, Christos J Petropoulos, and Javier Martinez-Picado. 2020. “Pan-Resistant HIV-1 Emergence in the Era of Integrase Strand-Transfer Inhibitors: A Case Report.” The Lancet Microbe 1 (3): e130–35. https://doi.org/10.1016/S2666-5247(20)30006-9.

48. Qi, Li, Craig E. Higgins, Stephen P. Higgins, Brian K. Law, Tessa M. Simone, and Paul J. Higgins. 2014. “The Basic Helix-Loop-Helix/Leucine Zipper Transcription Factor USF2 Integrates Serum-Induced PAI-1 Expression and Keratinocyte Growth: THE BHLH-LZ TRANSCRIPTION FACTOR USF2.” Journal of Cellular Biochemistry 115 (10): 1840–47. https://doi.org/10.1002/jcb.24861.

49. Qyang, Yibing, Xu Luo, Tao Lu, Preeti M. Ismail, Dmitry Krylov, Charles Vinson, and Michèle Sawadogo. 1999. “Cell-Type-Dependent Activity of the Ubiquitous Transcription Factor USF in Cellular Proliferation and Transcriptional Activation.” Molecular and Cellular Biology 19 (2): 1508–17. https://doi.org/10.1128/MCB.19.2.1508.

50. Rada-Iglesias, Alvaro, Adam Ameur, Philipp Kapranov, Stefan Enroth, Jan Komorowski, Thomas R. Gingeras, and Claes Wadelius. 2008. “Whole-Genome Maps of USF1 and USF2 Binding and Histone H3 Acetylation Reveal New Aspects of Promoter Structure and Candidate Genes for Common Human Disorders.” Genome Research 18 (3): 380–92. https://doi.org/10.1101/gr.6880908.

51. Roy, A. L. 1997. “Cloning of an Inr- and E-Box-Binding Protein, TFII-I, That Interacts Physically and Functionally with USF1.” The EMBO Journal 16 (23): 7091–7104. https://doi.org/10.1093/emboj/16.23.7091.

52. Sadowski, Ivan, and Farhad B. Hashemi. 2019. “Strategies to Eradicate HIV from Infected Patients: Elimination of Latent Provirus Reservoirs.” *Cellular and Molecular Life Sciences*, May. https://doi.org/10.1007/s00018-019-03156-8.

53. Sadowski, Ivan, Pedro Lourenco, and Tom Malcolm. 2008. “Factors Controlling Chromatin Organization and Nucleosome Positioning for Establishment and Maintenance of HIV Latency.” Current HIV Research 6 (4): 286–95. https://doi.org/10.2174/157016208785132563.

54. Sadowski, Ivan, and David A. Mitchell. 2005. “TFII-I and USF (RBF-2) Regulate Ras/MAPK- Responsive HIV-1 Transcription in T Cells.” European Journal of Cancer 41 (16): 2528–36. https://doi.org/10.1016/j.ejca.2005.08.011.

55. Sawadogo, M. 1985. “Interaction of a Gene-Specific Transcription Factor with the Adenovirus Major Late Promoter Upstream of the TATA Box Region.” Cell 43 (1): 165–75. https://doi.org/10.1016/0092-8674(85)90021-2.

56. Sawadogo, M., M. W. Van Dyke, P. D. Gregor, and R. G. Roeder. 1988. “Multiple Forms of the Human Gene-Specific Transcription Factor USF. I. Complete Purification and Identification of USF from HeLa Cell Nuclei.” The Journal of Biological Chemistry 263 (24): 11985–93.

57. Sayasith, Khampoune, Jacques G. Lussier, and Jean Sirois. 2005. “Role of Upstream Stimulatory Factor Phosphorylation in the Regulation of the Prostaglandin G/H Synthase-2 Promoter in Granulosa Cells.” Journal of Biological Chemistry 280 (32): 28885–93. https://doi.org/10.1074/jbc.M413434200.

58. Segura-Puimedon, Maria, Cristina Borralleras, Luis A. Pérez-Jurado, and Victoria Campuzano. 2013. “TFII-I Regulates Target Genes in the PI-3K and TGF-β Signaling Pathways through a Novel DNA Binding Motif.” Gene 527 (2): 529–36. https://doi.org/10.1016/j.gene.2013.06.050.

59. Shieh, B. H., R. S. Sparkes, R. B. Gaynor, and A. J. Lusis. 1993. “Localization of the Gene- Encoding Upstream Stimulatory Factor (USF) to Human Chromosome 1q22-Q23.” Genomics 16(1): 266–68. https://doi.org/10.1006/geno.1993.1174.

60. Sieweke, M. H. 1998. “Cooperative Interaction of Ets-1 with USF-1 Required for HIV-1 Enhancer Activity in T Cells.” The EMBO Journal 17 (6): 1728–39. https://doi.org/10.1093/emboj/17.6.1728.

61. Sirito, Mario, Qun Lin, Jian Min Deng, Richard R. Behringer, and Michèle Sawadogo. 1998. “Overlapping Roles and Asymmetrical Cross-Regulation of the USF Proteins in Mice.” Proceedings of the National Academy of Sciences 95 (7): 3758–63. https://doi.org/10.1073/pnas.95.7.3758.

62. Sirito, Mario, Qun Lin, Tapati Maity, and Michèle Sawadogo. 1994. “Ubiquitous Expression of the 43- and 44-KDa Forms of Transcription Factor USF in Mammalian Cells.” Nucleic Acids Research 22 (3): 427–33. https://doi.org/10.1093/nar/22.3.427.

63. Spohrer, Sarah, Rebecca Groß, Lisa Nalbach, Lisa Schwind, Heike Stumpf, Michael D. Menger, Emmanuel Ampofo, Mathias Montenarh, and Claudia Götz. 2017. “Functional Interplay between the Transcription Factors USF1 and PDX-1 and Protein Kinase CK2 in Pancreatic β-Cells.” Scientific Reports 7 (1): 16367. https://doi.org/10.1038/s41598-017-16590-0.

64. Swiggard, William J, Clifford Baytop, Jianqing J Yu, Jihong Dai, Chuanzhao Li, Richard Schretzenmair, Ted Theodosopoulos, and Una O’Doherty. 2005. “Human Immunodeficiency Virus Type 1 Can Establish Latent Infection in Resting CD4L T Cells in the Absence of Activating Stimuli.” J. VIROL. 79: 10.

65. Temereanca, Aura, and Simona Ruta. 2023. “Strategies to Overcome HIV Drug Resistance- Current and Future Perspectives.” Frontiers in Microbiology 14 (February): 1133407. https://doi.org/10.3389/fmicb.2023.1133407.

66. Terragni, Jolyon, Gauri Nayak, Swati Banerjee, Jose-Luis Medrano, Julie R. Graham, James F. Brennan, Sean Sepulveda, and Geoffrey M. Cooper. 2011. “The E-Box Binding Factors Max/Mnt, MITF, and USF1 Act Coordinately with FoxO to Regulate Expression of Proapoptotic and Cell Cycle Control Genes by Phosphatidylinositol 3-Kinase/Akt/Glycogen Synthase Kinase 3 Signaling.” Journal of Biological Chemistry 286 (42): 36215–27. https://doi.org/10.1074/jbc.M111.246116.

67. Tiruppathi, Chinnaswamy, Dheeraj Soni, Dong-Mei Wang, Jiaping Xue, Vandana Singh, Prabhakar B Thippegowda, Bopaiah P Cheppudira, et al. 2014. “The Transcription Factor DREAM Represses the Deubiquitinase A20 and Mediates Inflammation.” Nature Immunology 15 (3): 239–47. https://doi.org/10.1038/ni.2823.

68. Tong, Ann-Jay, Xin Liu, Brandon J. Thomas, Michelle M. Lissner, Mairead R. Baker, Madhavi D. Senagolage, Amanda L. Allred, Grant D. Barish, and Stephen T. Smale. 2016. “A Stringent Systems Approach Uncovers Gene-Specific Mechanisms Regulating Inflammation.” Cell 165(1): 165–79. https://doi.org/10.1016/j.cell.2016.01.020.

69. UNAIDS. 2022. “In Danger — UNAIDS Global AIDS Update 2022,” 376.

70. Unutmaz, Derya, Vineet N. KewalRamani, Shana Marmon, and Dan R. Littman. 1999. “Cytokine Signals Are Sufficient for HIV-1 Infection of Resting Human T Lymphocytes.” Journal of Experimental Medicine 189 (11): 1735–46. https://doi.org/10.1084/jem.189.11.1735.

71. Viollet, Benoît, Anne-Marie Lefrançois-Martinez, Alexandra Henrion, Axel Kahn, Michel Raymondjean, and Antoine Martinez. 1996. “Immunochemical Characterization and Transacting Properties of Upstream Stimulatory Factor Isoforms.” Journal of Biological Chemistry 271 (3): 1405–15. https://doi.org/10.1074/jbc.271.3.1405.

72. Walker-Sperling, Victoria E., Christopher W. Pohlmeyer, Patrick M. Tarwater, and Joel N. Blankson. 2016. “The Effect of Latency Reversal Agents on Primary CD8 + T Cells: Implications for Shock and Kill Strategies for Human Immunodeficiency Virus Eradication.” EBioMedicine 8 (June): 217–29. https://doi.org/10.1016/j.ebiom.2016.04.019.

73. Wang, Yuhui, Roger H.F. Wong, Tianyi Tang, Carolyn S. Hudak, Di Yang, Robin E. Duncan, and Hei Sook Sul. 2013. “Phosphorylation and Recruitment of BAF60c in Chromatin Remodeling for Lipogenesis in Response to Insulin.” Molecular Cell 49 (2): 283–97. https://doi.org/10.1016/j.molcel.2012.10.028.

74. West, Adam G., Suming Huang, Miklos Gaszner, Michael D. Litt, and Gary Felsenfeld. 2004. “Recruitment of Histone Modifications by USF Proteins at a Vertebrate Barrier Element.” Molecular Cell 16 (3): 453–63. https://doi.org/10.1016/j.molcel.2004.10.005.

75. Wong, Roger H. F., and Hei Sook Sul. 2009. “DNA-PK: Relaying the Insulin Signal to USF in Lipogenesis.” Cell Cycle 8 (13): 1973–78. https://doi.org/10.4161/cc.8.13.8941.

76. Xiao, Qianxun, Agnes Kenessey, and Kaie Ojamaa. 2002. “Role of USF1 Phosphorylation on Cardiac α-Myosin Heavy Chain Promoter Activity.” American Journal of Physiology-Heart and Circulatory Physiology 283 (1): H213–19. https://doi.org/10.1152/ajpheart.01085.2001.

77. Zhang, Hao, Yang Zhang, Xinyue Zhou, Shaela Wright, Judith Hyle, Lianzhong Zhao, Jie An, et al. 2020. “Functional Interrogation of HOXA9 Regulome in MLLr Leukemia via Reporter-Based CRISPR/Cas9 Screen.” ELife 9(October): e57858. https://doi.org/10.7554/eLife.57858.

