## Supplementary Information for "Upstream Stimulatory Factors regulate HIV-1 latency and are required for robust T cell activation"

#### Legends to Supplementary Figures

**Figure S1. shRNA knockdown of *USF2* inhibits HIV-1 expression. Panels A, C:** Jurkat mHIV-Luciferase cells were transduced with an pLKO empty vector control or shRNA targeting USF1 (A) or USF2 (C). Transduced cells were selected with 3 μg/mL puromycin for up to 8 days after which lysates were analyzed by immunoblotting using antibodies targeting USF1, USF2, and Tubulin. **Panel B, D:** USF1 (B) or USF2 (D) shRNA transduced mHIV-Luciferase cells were left untreated (Ve, DMSO) or stimulated with 5 nM PMA for 4 hours prior to measuring luciferase activity ($n$ = 3, mean ± SD).

**Figure S2. Expression of USF1 or USF2 rescues HIV-1 transcription in *USF2* KO cells. Panel A:** Schematic representation of the LAI LTR-Tat-IRES-GFP reporter construct and the CMV-EV-EF1α-RFP (EV), CMV-USF1-EF1α-RFP (USF1), and CMV-USF2-EF1α-RFP (USF2) expression vectors. **Panel B:** Flow cytometry scatter plot generated following co-transfection of LTR reporter and RFP expression vector. The bottom left quadrant (Q4) contains un-transfected cells, negative fluorescent cells. The top left population (Q1) is co-transfected by the LTR reporter and the RFP expression vector does not express the LTR reporter. Cells in the top right quadrant (Q2) are co-transfected by transcriptionally active LTR reporter as well as the RFP expression vector. The bottom right quadrant (Q3) are cells transfected only by the active LTR reporter. **Panel C:** Representative FACS scatter plots following co-transfection of wildtype or *USF2* KO HEK293T cells with the LTR-Tat-IRES-GFP and either CMV-EV-EF1α-RFP, CMV-USF1-EF1α-RFP (USF1), or CMV-USF2-EF1α-RFP (USF2) vector.

**Figure S3. USF2 regulates HIV expression independently of Tat. Panel A:** Schematic representation of the CMV-EV-EF1α-RFP expression vector and LAI LTR-IRES-GFP (-Tat) and LTR-Tat-IRES-GFP (+Tat) reporter. **Panel B:** Depiction of the flow cytometry scatter plot generated following co-transfection of LTR reporter and RFP expression vector. See Fig. S3*B* for further description. **Panels C, D:** Representative scatter plots following co-transfection of wildtype or *USF2* KO HEK293T cell lines with RFP expression vector and either LTR-IRES-GFP (C) or LTR-Tat-IRES-GFP (D) LAI derived reporter construct.

**Figure S4. Loss of USF1 or USF2 impairs expression of genes downstream of multiple T cell activation pathways. Panels A - D:** Hallmark GSEA plots of RNA-seq data analyzed by DESeq2 of PMA/ionomycin stimulated *USF1* KO and *USF2* KO Jurkat T cells. Analysis was performed on $n = 3$ RNA-seq samples.

**Figure S5. Representative flow cytometry gating strategy. Panels A, B:** Representative scatter plots depicting the gating strategy employed for HEK293T cell lines (A) and Jurkat T cell lines (B). Homogeneous populations of live cells were isolated and assessed by setting threshold forward scatter (FSC) and side scatter (SSC) settings.

### Supplementary Figure 1

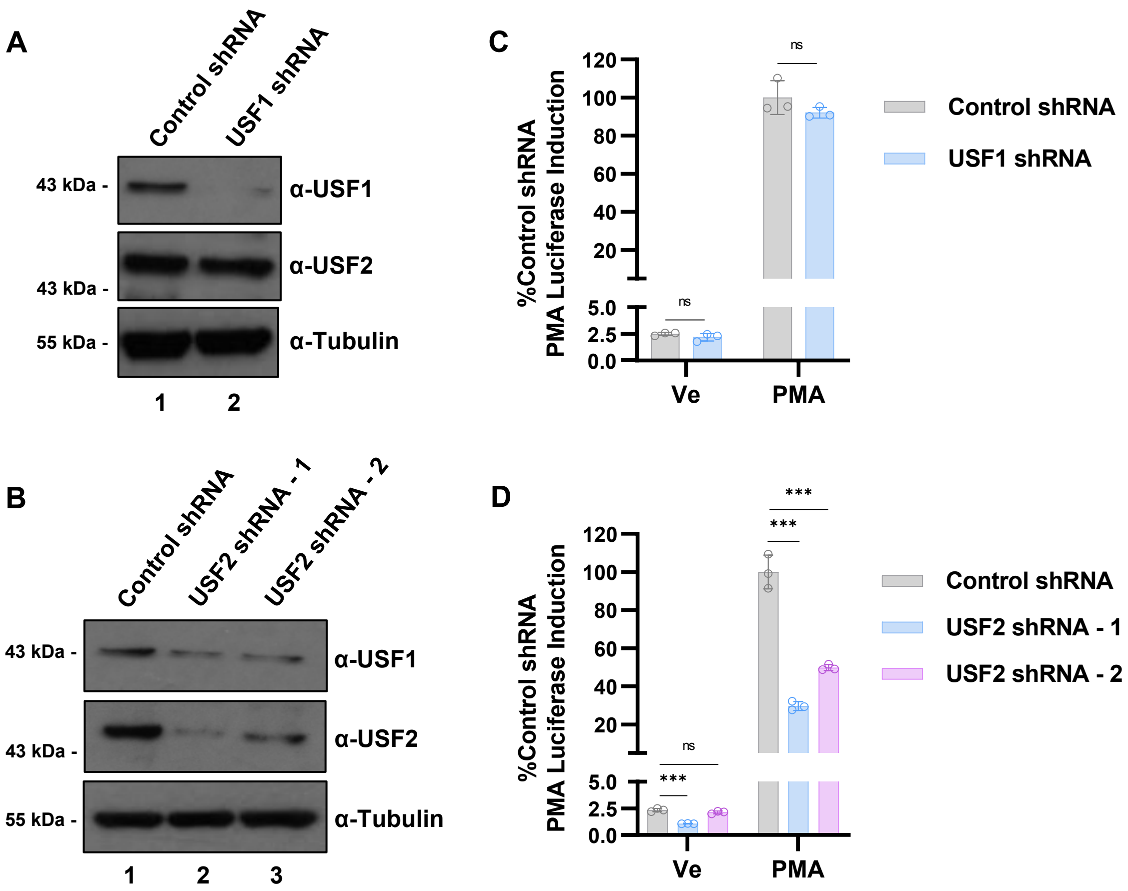

### Supplementary Figure 2

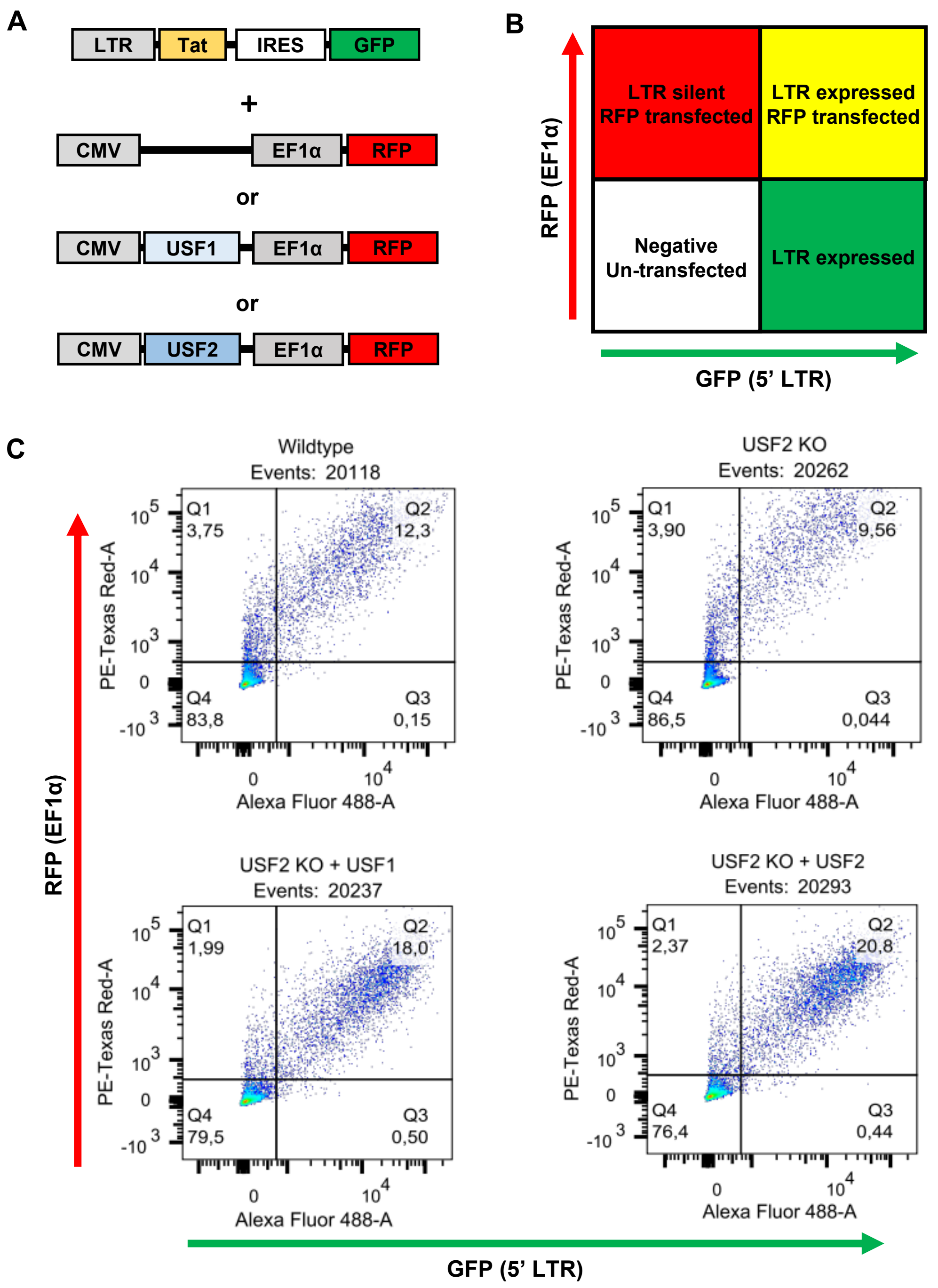

### Supplementary Figure 3

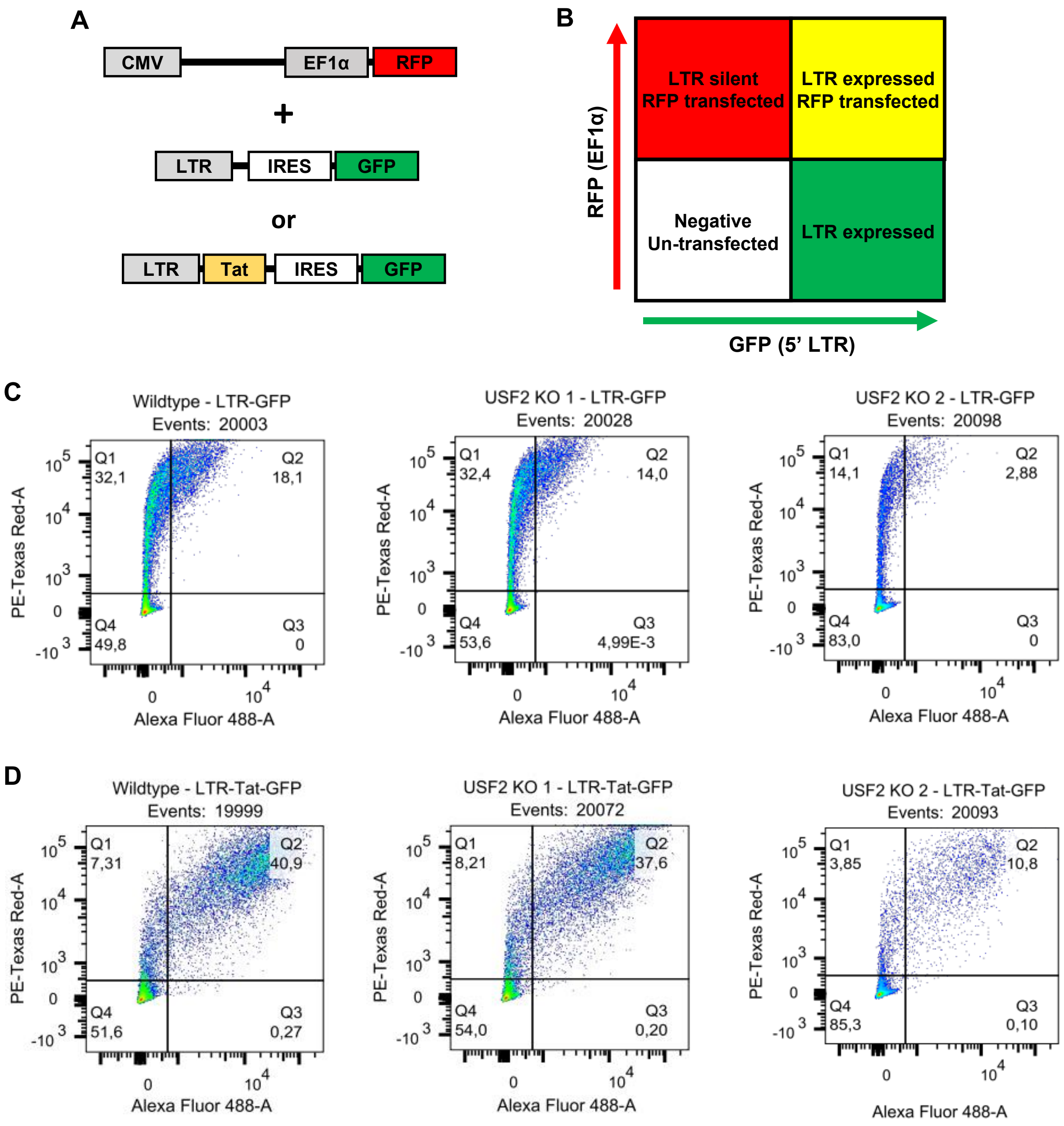

### Supplementary Figure 4

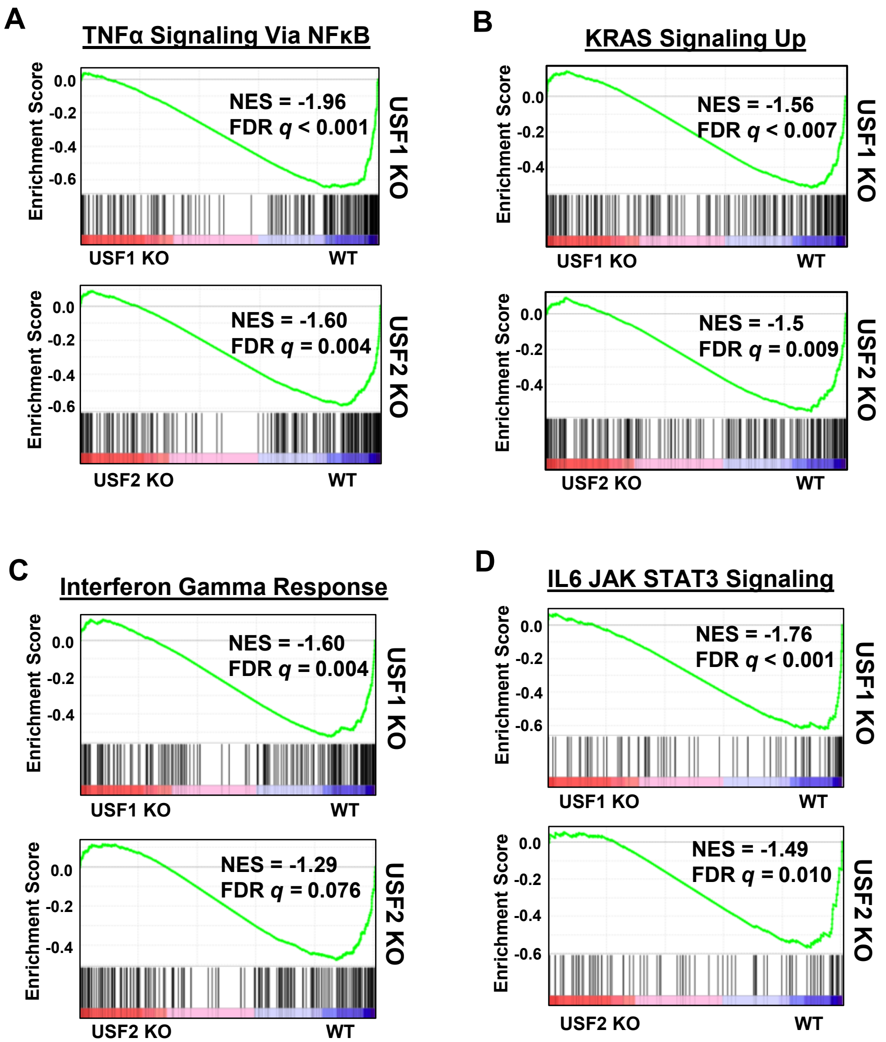

### Supplementary Figure 5

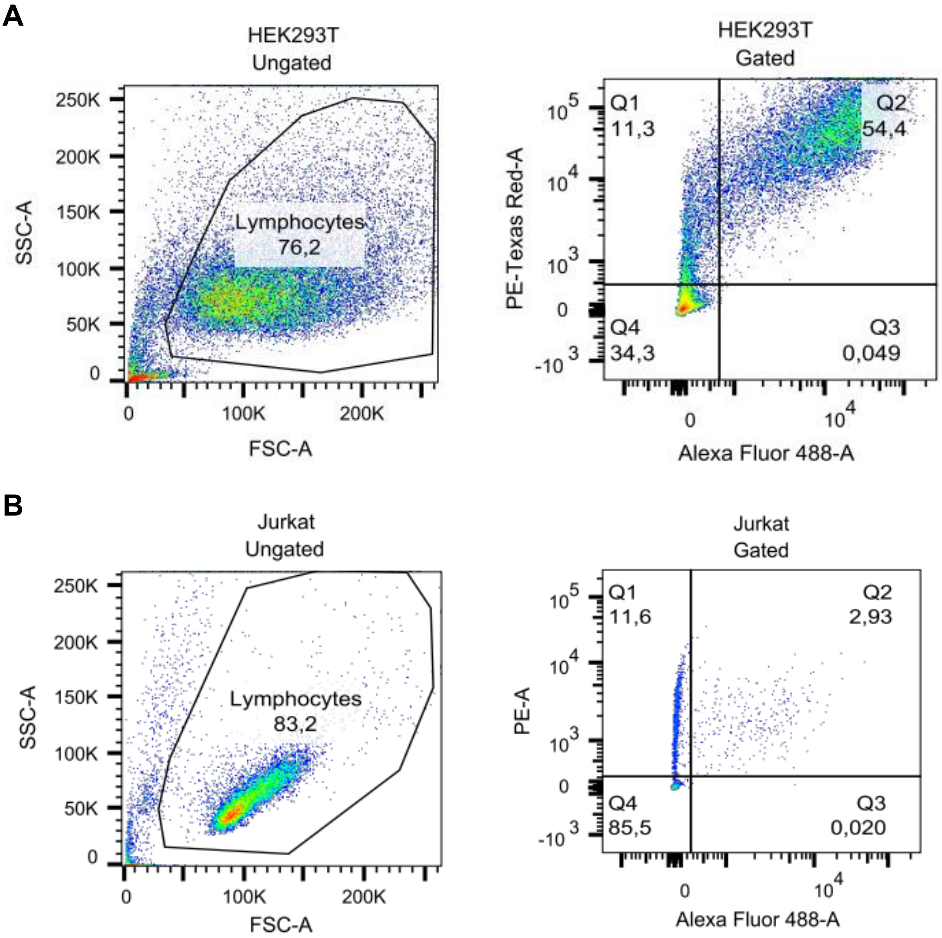
